# Demographic history, cold adaptation, and recent NRAP recurrent convergent evolution at amino acid residue 100 in the world northernmost cattle from Russia

**DOI:** 10.1101/2020.06.15.151894

**Authors:** Laura Buggiotti, Andrey A. Yurchenko, Nikolay S. Yudin, Christy J. Vander Jagt, Hans D. Daetwyler, Denis M. Larkin

## Abstract

Native cattle breeds represent an important cultural heritage. They are a reservoir of genetic variation useful for properly responding to agriculture needs in light of ongoing climate changes. Evolutionary processes that occur in response to extreme environmental conditions could also be better understood using adapted local populations. Herein, different evolutionary histories for two of the world northernmost native cattle breeds from Russia were investigated. They highlighted Kholmogory as a typical taurine cattle, while Yakut cattle separated from European taurines ~5,000 years ago and contain numerous ancestral and some novel genetic variants allowing their adaptation to harsh conditions of living above the Polar Circle. Scans for selection signatures pointed to several common gene pathways related to adaptation to harsh climates in both breeds. But genes affected by selection from these pathways were mostly different. A Yakut cattle breed-specific missense mutation, H100Q, in a highly conserved *NRAP* gene, represents a unique example of a young amino acid residue convergent change shared with at least 16 species of hibernating/cold-adapted mammals from nine distinct phylogenetic orders. This suggests a convergent evolution event along the mammalian phylogenetic tree and fast fixation in a single isolated cattle population exposed to a harsh climate.

## Introduction

Cattle domestication occurred ~8,000–10,000 years ago as a result of at least two independent events in the Fertile Crescent and the Indus Valley from two main sources of ancient *Bos* subspecies, *B. taurus* and *B. indicus* (1, 2). Many contemporary breeds adapted to a vast variety of environmental conditions originate from ancient and/or recent interbreeding between the *B. taurus* (taurine) and *B. indicus* (indicine) populations (2) and admixture with several other bovinae species, including yak, banteng, and gaur (3, 4). While adaptations of cattle breeds to hot climates are relatively well studied due to the economic needs in light of global warming (5–7), there is limited knowledge about the mechanisms of cattle adaptation to colder climates with most studies limited to genotyping datasets (8) rather than whole genome (re)sequencing. There are several northern cattle breeds that are of a particular interest for such studies. Two of them originate from, and are adapted to, the harsh climates of Russia: the Kholmogory and the Yakut cattle. The Kholmogory cattle breed was formed in the European part of Russia (Arkhangelsk District), about 300 years ago, from local taurine landraces which were crossed with an imported ‘Dutch cattle’ as early as in the 18^th^ century (9). Nowadays, the Kholmogory cattle is a highly productive dairy breed, which can be found in different parts of Russia including the northern territories (Siberia). The Yakut cattle (*Sakha Ynaga*) have a different history. They were likely formed at the Baikal area of Siberia (10) and migrated together with Yakut people to the contemporarily Yakutia region about 500-800 years ago (10). They now are the world northernmost cattle population which can be found up to 200 km above the Polar Circle, where they are exposed to long winters and low temperatures that occasionally can drop to below −70°C. At the territory of their original formation they were likely influenced by Asian cattle populations and belong therefore to the so-called ‘Turano-Mongolian’ cattle group. Yakut cattle represent the only extant pure commercial population from this group as most other Turano-Mongolian breeds were interbred with European taurines to improve performance (9). Indeed, a recent phylogenetic analysis placed the Yakut cattle together with other Turano-Mongolian breeds, including the Hanwoo and Japanese Black, but at the same time suggested possible historical admixture with indicine and African taurine cattle (11). However, the origin of this Turano-Mongolian cattle group that is highly divergent from the European taurines remains unclear with some authors suggesting an independent domestication event in Eastern Asia (12, 13).

Studies of multiple cold adapted species show that they can temporarily slow their metabolisms in response to harsh environmental conditions, leading to torpor or, in extreme cases, to hibernation (14). Ability/inability to hibernate could be observed in species from the same phylogenetic node (e.g. there are hibernating and non-hibernating bats) suggesting that this could be an example of convergently evolving phenotype (15, 16). Other researches, however, propose that the ability to hibernate could be an ancestral mechanism lost in some lineages (17). Comparison of selected genes in multiple Arctic and Antarctic mammals pointed to a set of genes that could be independently selected in response to cold adaptation. These include genes involved in hypoxia, actin binding, extracellular matrix etc., highlighting common pathways of cold adaptation (18). The study of Arctic and Antarctic fishes provides an example of independent changes in antifreeze glycoproteins allowing these species to survive sub-zero temperatures (19). In Antarctic fishes the antifreeze protein evolved from trypsinogen-like serine protease gene, while the Arctic polar cod genome contains nearly identical proteins evolved from a different genomic locus, suggesting convergent evolution (19). A recent study of Russian cattle and sheep breed genomes (20) found putative selection in regions containing known candidate genes for cold adaptations (21), but failed to identify if there could be any convergent molecular changes in the recent evolution of artificially designed livestock populations because of the limited data resolution (~100,000 SNP loci in the genome). Previous studies have indicated that convergent amino acid changes contribute to appearance of similar features in phylogenetically distinct groups of animals (22), environmental and ecological adaptations (23, 24), but this phenomenon was not so far found in individual breeds of domesticated species.

The present study focuses on comparison of resequenced genomes from two cold-adapted cattle breeds from Russia, the Kholmogory and Yakut cattle. These distinct populations were compared to their phylogenetically closest breeds to identify signatures of selection and copy number variants that could have influence on their biology. In addition, the demographic and admixture analysis of the two populations suggests that the Kholmogory and Yakut cattle indeed belong to two distinct branches of taurine cattle, which became separated about 4,000-5,000 years ago. Our data suggest that while the Kholmogory cattle are a typical European taurine cattle breed, the Yakut cattle genome contains a distinct fraction of a largely ancestral cattle variants, either being introgressed into the Yakut cattle genome or being lost in the European taurines. The two breeds demonstrate signatures of selection in the same gene pathways related to adaptation to harsh climates, but the genes affected by selection from these pathways are mostly different. All resequenced Yakut cattle individuals contain a missense mutation in a highly conserved *NRAP* gene. This change is shared with a majority of hibernating mammals but absent from all other cattle breeds tested and a majority of mammalian species, suggesting a recent convergent evolution event and fast selection in a single isolated cattle population exposed to a harsh climate.

## Materials and Methods

### Sample selection and genome sequencing

Twenty samples per breed from the *Yakut* and *Kholmogory* Russian native cattle breeds were selected based on their kinship coefficient (to exclude closely related animals) and maximum purity estimated by the plink software command –genome and ADMIXTURE software respectively from our previous genotyping of the same samples (20). Next generation sequencing (NGS) paired end reads (PE, 150bp) were generated using HiSeq sequencing platform at Novogene Co., Ltd. Hong Kong, China. Library preparation was carried out by Novogene and raw sequence data per sample, ranging from 41.1 to 66.8 Gb, were generated. In addition, we obtained raw sequence data of 153 cattle individuals from the NCBI SRA database, including 20 Holstein (25), 20 Hanwoo (26), 47 African cattle (Ankole – ten, Boran – nine, Kenana – nine, nDama – ten, Ogaden – nine), 36 Chinese cattle (Luxi – five, Nanyang – five, JaxianRed – five, Dianzhong – six, Wannan – five, Ji’an – four, Leiqiong – three, Yanbian - three), five Brahman, ten Yak (*Bos grunniens*), and 15 of other *Bos* species (five Gayal (*Bos frontalis*), five Banteng (*Bos javanicus*), three Gaur (*Bos gaurus*), two Bison (*Bison bison*); Supplementary Information 1) (27, 28). Reference Hereford cattle (*Bos taurus*) assembly (UMD3.1, *BosTau6*) was downloaded from NCBI.

### Read mapping, and variant calling

All cleaned reads were mapped to the cattle reference assembly using BWA-MEM (29) with default parameters. The average mapping rate of the reads generated in this study was 99.68%, and the average sequencing coverage was approximately 11.43X (ranging from 9.7 to 15.1, Table 1). Alignment preprocessing steps and variant calling were done following the Genome Analysis Toolkit (GATK v. 3.8 (30)) pipeline. For each raw BAM file, we marked duplicate reads with Picard (v. 1.69) using the tool MarkDuplicates (http://broadinstitute.github.io/picard/). Next, we performed base quality score recalibration (using cattle known variants: dbSNP148). Variant calling was performed using the Haplotype Caller tool from GATK. Three joint VCF files were generated for: 1) the Yakut, Kholmogory, Holstein, and Hanwoo breeds, 2) the Yakut, Kholmogory, Holstein, Hanwoo, Indian, Chinese and African breeds, and 3) the Yakut, Kholmogory, Holstein, Hanwoo, Indian, Chinese and African breeds and bovinae species samples (see Supplementary Information 1 for samples). The filtering of SNPs for quality (“hard filtering”) was applied with the following parameters: i) variant confidence/quality by depth <2; ii) RMS mapping quality (MQ)<40.0; iii) Phred-scaled p-value using Fisher’s exact test to detect strand bias > 60; iv) Z-score from the Wilcoxon rank sum test of alternative vs. reference read MQs (MQRankSum) < −12.5; and (v) Z-score from the Wilcoxon rank sum test of alternative vs. reference read position bias (ReadPosRankSum)<−8. The thresholds for these parameters were adopted from GATK Best Practices (30). Multi-allelic SNPs and INDELs were removed and the final filtered VCF file was used for further analysis.

**Table 1.** Summary of samples and genome sequencing statistics (hard filtered biallelic SNP).

|  | Holstein | Kholmogory | Hanwoo | Yakut | Total |
| --- | --- | --- | --- | --- | --- |
| **Samples** | 20 | 20 | 20 | 20 | 80 |
| **Coverage** | 10.60±5.1 | 11.68±1.2 | 10.21±3.13 | 11.18±0.76 |  |
| **SNPs** | 14,448,782 | 15,426,893 | 18,914,454 | 17,025,030 | 27,487,145 |
| **ts/tv** | 2.13 | 2.20 | 2.23 | 2.23 | 2.15 |
| **Synonymous\*** | 56,249 | 49,976 | 56,568 | 75,798 | 109,774 |
| **Missense\*** | 32,216 | 31,089 | 35,740 | 35,919 | 64,221 |
| **CDS** | 53,523 | 63,551 | 70,590 | 92,202 | 137,744 |
| **3' UTR** | 33,537 | 36,100 | 45,514 | 46,886 | 74,332 |
| **5' UTR** | 6,345 | 7,419 | 8,473 | 8,557 | 13,814 |
| **Intron** | 2,927,686 | 3,100,776 | 3,898,489 | 3,457,265 | 5,676,380 |
| **Intergenic** | 11,489,562 | 12,280,607 | 14,965,760 | 13,485,886 | 21,685,165 |
| **MamCNEs** | 82,533 | 85,465 | 108,525 | 89,593 | 167,632 |
| **CetCNEs** | 42,643 | 43,811 | 56,327 | 45,934 | 84,970 |
| **RumCNEs** | 74,576 | 76,886 | 100,8716 | 83,653 | 153,684 |

\*Synonymous and missense mutations were annotated by dNdScv. Abbreviations are as follows: CNEs (conserved non-coding elements); MamCNEs (Mammalian CNEs); CetCNEs (Cetartiodactyl CNEs); RumCNEs (Ruminant CNEs).

### Genomic annotation of SNPs

Whole genome hard filtered SNPs were annotated using the Ensembl Variant Effect Predictor (VEP) with Ensembl v94 (31). SNPs were annotated with the classification categories of impact HIGH, LOW, MODERATE, and MODIFIER (Supplementary Information 2). Moreover, dNdScv (32), used to estimate whole genome dN/dS, also classified SNPs based on the RefSeq Release 103 (33).

We derived a subset of SNPs from the four breeds’ VCF files with the following characteristics: 1) high frequency of alternative (non-reference) allele in Yakut cattle (Yakut alternative allele frequency of >= 70% and <=10% in Holstein, Kholmogory, and Hanwoo), 2) SNPs present in at least 25% of the four breed sample set. NGS annotation pipeline (34) was used to assign classification (e.g., intronic, missense, synonymous, splice variant, stop gain, etc.) to each SNP in this subset of SNPs and to provide several fields of information describing the affected transcript and protein, if applicable. NGS-SNP also reports NCBI, Ensembl, and UniProt IDs for genes, transcripts, and proteins when applicable. For missense SNPs, an “alignment score change” was calculated by comparing the reference amino acid and the non-reference amino acid to an orthologous amino acid in a range of genomes. A positive value indicated that the alternative allele amino acid was more common in the orthologous position in other species than the reference amino acid, whereas a negative value indicated that the reference amino acid was more common (Supplementary Information 3).

### Demographic history reconstruction

To calculate historical demography and separation time between pairs of populations we used two complementary approaches: SMC++ (35) and GADMA (36), which use different population inference models. To remove spurious variants and regions with bad mapping quality from further consideration, we first computed the mappability score of the bovine reference genome using the GEM mappability program (37) with kmer size =100 bp allowing one mismatch. The regions with mappability < 1 were removed. Additionally, we removed all regions under selection identified in the bovine genome (38) and sex chromosomes.

SMC++ uses a sequential Markovian Coalescent approach to infer demographic history of separate populations and pairwise time of divergence between them on the unphased samples. The drawback of this method is an inability to implement migration into the model. For the SMC++ we analyzed demographic history using the cubic spline approach with mutation rate per bp per generation = 1.2e-8.

The approach implemented in the GADMA software is based on the comparisons of observed and modeled joint allele frequency spectrums of populations (39, 40) and allows the efficient modelling of demographic parameters and migration, given a divergence scenario with up to three populations, using a powerful genetic algorithm. We ran GADMA with five years as cattle generation time using 10 processes simultaneously and linked loci. The main algorithm chosen was moments (ordinary differential equations).

### Detection of genetic introgression between breeds

We applied a robust forward-backward algorithm implemented in RFMix (41) to screen for the presence of putative taurine or indicine haplotypes in autosomes of the Yakut cattle. This algorithm uses designated reference haplotypes to infer local ancestry in designated admixed haplotypes, thus five genetic groups were selected as a reference panel: European taurine (Holstein), Russian taurine (Kholmogory), African indicine (Ogaden), Chinese indicine (Wannan, Ji’an, and Leiquiong), and Indian indicine (Brahman). Window size was set to three (-w 3) and the option “--reanalyze-reference” with three iterations was used to analyze the reference haplotypes as if they were query haplotypes (41).

We conducted TreeMix v1.12 (42) analyses to infer the relationships, divergence, and major mixtures among 18 cattle breeds and five bovinae species (Supplementary Information 1). We applied the option “-root YAK”, which sets Yak as the position of the root, option “-k 1000”, which builds the tree using blocks of 1000 SNPs to account for linkage disequilibrium and used the option “-se” to calculate standard errors of migration proportions. We allowed up to 15 migration edges on the tree (m ranging from 0 to 15) and generated a residual heatmap to identify populations that were not well-modeled after adding each migration edge. The percentages of variation explained by the maximum likelihood trees were also calculated. Migration edges were considered until 99.9% of the variance in ancestry between populations was explained by the model. We also ensured that the incorporated migration edges were statistically significant. Residuals from the model were visualized using the R script implemented in TreeMix. Finally, in order to provide further support for a past admixture between populations, we calculated *f3* and *D* statistics using ADMIXTOOLS (v 5.1) with default parameters (43). We calculated *f3* statistics using qp3Pop of the form (X; A, B) where a negative value mean implies that X is admixed with populations close to A and B. We considered negative statistics with Z-score values below −2 as significant signals of admixture.

We computed *D* statistics using qpDstat. *D* statistics of the form D (A, B, X, Y) to test the null hypothesis of the unrooted tree topology ((A, B), (X, Y)) was used. A positive value indicates that either A and X, or B and Y share more drift than expected under the null hypothesis. We quote *D* statistics as the Z score computed using default block jackknife parameters.

### Detection of copy number variants (CNVs) and copy number variant regions (CNVRs)

cn.MOPS (Copy number estimation by a Mixture Of PoissonS; (44)) was used for CNV region detection in four cattle breeds (Kholmogory, Yakut, Hanwoo, and Holstein) and pure indicine breeds (Brahman, Ogaden, Wannan, Ji’an, and Leiquiong) chosen based on the structure analysis reported by Chen et al (28). A total of 75 samples of the four cattle breeds were used for CNV detection, and five samples (one from Holstein, two from Hanwoo, and two from Yakut) were removed from the analysis as they were problematic probably due to their lower sequence coverage. Based on the average sequence coverage of our data, window length was set to 700 and posterior probabilities estimated. The tool represents a CNV detection pipeline that models the depths of coverage across multiple samples at each genomic position. Using a Bayesian approach, it decomposes read variations across samples into integer CNVs and noise using mixture components and Poisson distributions, respectively. The multiple samples approach increases statistical power and decreases computational burden and the FDR in CNV detection. CNVs were then used to construct a set of copy number variable regions (CNVRs) for Kholmogory, Yakut, Hanwoo, and Holstein. Moreover, we applied the following filtration steps: i) cn.MOPs classes one, two, and three (CN1-CN3) were considered as normal and the rest (CN0, CN4-CN8) were considered as CNVs; ii) CNVRs were constructed by merging CNVs across samples of the same breed that exhibited at least 50% pairwise reciprocal overlap in their genomic coordinates; iii) unique CNVRs per breed were those having less than 10% overlap with CNVRs in other breeds. In addition, cn.MOPS was run a second time by using the four above mentioned breeds together with additional 26 individuals of Indian (5), Chinese (12), and African (9) indicine origin, to detect indicine CNVs. BEDTools (45) and BEDOPS (46) tools were used to calculate CNVR overlaps.

### Genome scan for selection signals

#### Identification of signatures of selection with hapFLK statistics

We performed a genome scan for selective sweeps within the groups of phylogenetically closely related breeds: Yakut and Hanwoo and Kholmogory and Holstein using haplotype-based statistics hapFLK (47). Due to the hapFLK model assuming that selection acts on shared ancestral SNP allele frequencies, we excluded rare SNPs with low minor allele frequencies (MAFs) from breed groups (MAF<0.1). We also excluded poorly genotyped individuals (<95% of SNPs with genotypes), loci genotyped in <99% of samples, SNPs without chromosomal assignments, and SNPs on sex chromosomes in PLINK, using the commands: --maf 0.1, --mind 0.05, --geno 0.01, and --chr 1-29 prior to performing the genome selection scans. This resulted in 8,423,043 and 7,926,039 SNPs for Yakut-Hanwoo and Kholmogory-Holstein groups, respectively.

The hapFLK method takes the haplotype structure of the population into account. What was important for our dataset is that this method can account for population bottlenecks and migration. Reynolds distances and a kinship matrix were calculated by the hapFLK program v.1.4 (47). For the hapFLK analysis, the number of haplotype clusters for each breed group were estimated with fastPhase (48) and were set as -*K* 10 for the Yakut-Hanwoo, and -*K* 20 for Kholmogory-Holstein groups. The expected maximum number of iterations was set to 30 for both groups. We applied midpoint rooting to all sets of breeds.

#### P-value calculation

For hapFLK, the calculation of raw p-values was performed assuming that the selected regions represent only a small fraction of the genome (49). The genome-wide distribution of hapFLK statistics could be modelled relatively well with a normal distribution except for a small fraction of outliers from potentially selected regions (49). Robust estimations of the mean and variance of the hapFLK statistic were obtained using the *R* MASS package *rlm* function to eliminate influence of outlying regions following Biotard and co-workers (50). This has been done for each group (Yakut-Hanwoo, and Kholmogory-Holstein). The hapFLK values were Z-transformed using these parameter estimates, and p-values were calculated from the normal distribution in *R*. The *R qvalue* package was used to correct p-values for multiple testing (51). To identify selected intervals, we scanned the hapFLK output for intervals containing q-values < 0.01. The boundaries of the selected intervals were defined by the first marker up- and down-stream with a q-value > 0.1. To assign a selection signal to a specific breed Local Reynolds distances were calculated for selected regions using *local_reynolds.py* script and local population trees were then built with the *local_trees* R script obtained from https://forge-dga.jouy.inra.fr/projects/hapflk/wiki/LocalTrees.

#### Identification of genomic regions under selection: dN/dS ratio

Whole genome dN/dS estimates were obtained for the SNPs found in any of the four cattle breeds (Yakut, Kholmogory, Holstein, and Hanwoo) using the dNdScv (32). The background mutation rate of each gene was estimated by combining local information (synonymous mutations in the gene) and global information (variation of the mutation rate across genes) and controlling for the sequence composition of the gene and mutational signatures. Unlike traditional implementations of dN/dS, dNdScv uses trinucleotide context-dependent substitution matrices to avoid common mutation biases affecting dN/dS (52). The calculations were initially made using all hard-filtered SNPs (MAF in each breed >= 0.1) using the *Btau6* reference genome database generated and the “dndscv” function. The second run was done using hard filtered SNPs with MAF>= 0.6 in a breed to identify genes under selection which likely affected the whole breed. One sided p-values were calculated and adjusted for multiple testing using Benjamini and Hochberg’s false discovery rate to detect both positive and negative selection (q-value positive selection and q-value negative selection).

### F*_ST_* calculations against the 1000 Bull Genome Dataset

Fixation index of genetic differentiation (F*_ST_*) between biallelic SNPs identified in our Yakut or Kholmogory cattle samples as part of their analysis by the 1000 Bull Genome Project Run 7 against the reference cattle ARS-USD1.2 genome was calculated against: a) the set of pure taurine breeds (2,003 animals, 18 breeds) as identified from the structure analysis reported by Chen et al (28) and b) against all cattle samples of taurine or indicine origins (3,362 animals) from the 1000 Bull Genome Project Run 7. This was performed using *vcftools* (53) for each cattle autosome applying the following settings: --*fst-window-size 10000* --*fst-window-step 5000*. All animals from the 1000 Bull Genome Project Run 7 have been filtered for a minimum of 6x mean genome coverage. One percent of the windows with the maximum mean F_*ST*_ values were extracted and overlapped with the gene set from Ensembl v.94.

### Functional enrichment analysis and protein modeling

Cattle genes based on the Ensembl v.87 database were retrieved using the Ensembl BioMart online tool (54) and gene ontology (GO) analysis was performed using genes found in unique CNVRs in each breed: ‘indicine-like’ genes from the RFMix analysis, and the hapFLK selected intervals using an in house pipeline. Moreover, a David functional cluster analysis (55) and a Reactome (56) pathway enrichment analysis were performed.

Genomic Association Test – GAT (57) was used to estimate the significance of overlap between multiple sets of genomic intervals by using 10K permutation. The genomic territory was represented by the *Btau6* whole genome and the tool was used to assess the significance of unique CNVRs among the four cattle breeds, and indicine-like assigned genes from the RFMix analysis.

Protein sequences for several species and their alignments were retrieved from the UniProt database (58) and ESPript v3.0 (59) was used to gather sequence similarities and secondary structure information from aligned sequences. Next, Phyre2 (60) allowed to predict protein 3D structure and estimate mutation sensitivity; the “intense” modelling option was used.

## Results

### Whole genome resequencing, mapping, and SNP detection

Resequencing of the Yakut and Kholmogory cattle samples resulted in a total of 1,667.61 Gb of filtered sequence data. The read mapping rate against the reference genome for Kholmogory cattle was 99.62%, while for the Yakut cattle it was 99.59%, resulting in an average sequencing coverage of 11.68x and 11.18x for the Kholmogory and Yakut samples, respectively (Table 1). The total number of SNPs was 17.03 and 15.43 million for the Yakut and Kholmogory samples, respectively. To identify copy number variants (CNVs) and signatures of selection differentiating the Yakut and Kholmogory cattle breeds from phylogenetically most related breeds (20), 20 samples of Holstein and 20 samples of Hanwoo breeds were included in our dataset. The average sequencing coverage per sample was 10.60x for Holsetin and 10.21x for Hanwoo. A total of 31.08 million SNPs were called in the four-breed joint set of which 27.49 million passed our quality filtering criteria. As expected, the majority (21.69 million (78.90%)) of the SNPs were located in the intergenic and intronic regions. About 0.60%, 0.31%, and 0.56% of the SNPs overlapped mammalian, cetartiodactyl, and ruminant conserved non-coding elements (61) and the number of synonymous and missense SNPs were 109,774 and 64,221, respectively (Table 1). There were 35,919 and 31,089 missense variants in the Russian Yakut and Kholmogory breeds, respectively. We found 224 and 210 nonsense variants in the Yakut and Kholmogory breeds, of which 32.60% and 21.90% were found in a single breed, respectively. A total of 678 and 594 SNPs were classified as “essential splice variants” while 17 and 15 were stop losses, in the Yakut and Kholmogory breeds, respectively (Supplementary Information 2).

To identify historical relationships and admixture between the Yakut cattle and other cattle breeds and species with potential historical admixture, SNPs from additional 113 individuals, representing indicine (African, Chinese, and Indian) cattle and five bovinae species (yak, banteng, gayal, gaur, bison) were combined with the four breed SNP set, resulting in a total of 193 individuals (Supplementary Information 1) and 11.03 million high quality biallelic SNPs.

### Demographic history and population divergence

We estimated demographic and population history using two methodologically distinct approaches: coalescent-based SMC++ (35) and ordinary differential equations (moments) realised in the GADMA package (36). To understand demographic histories of the Kholmogory and Yakut cattle breeds we focused to the three breeds with largest sample data and the least possibility of a recent admixture: Yakut, Kholmogory, and Holstein. SMC++ algorithm estimated a stepwise decline in the population size of cattle breeds with Indicine lineage (Brahman) as the most divergent and a much less pronounced recent decline. Yakut cattle demonstrated much stronger population decline more recently (around 200 – 500 years ago) than Kholmogory and Holstein (Fig. 1D).

**Figure 1.**
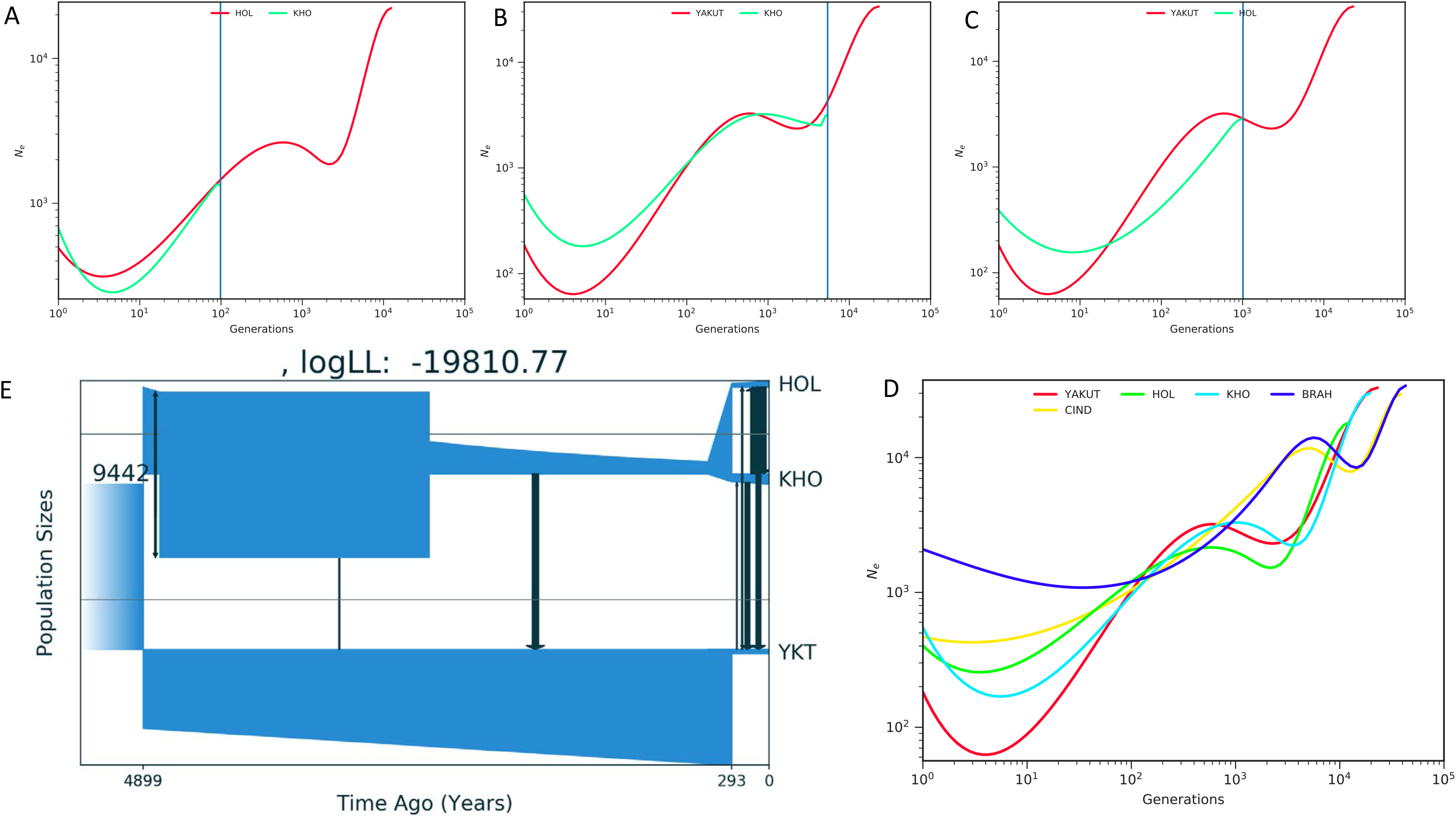
Demographic history of Yakut and Kholmogory cattle breeds. SMC++-inferred effective population sizes (Ne) with respect to time (generations) for A) Holstein (HOL) and Kholmogory (KHO), B) Yakut and KHO, C) Yakut and HOL, and D) Yakut, HOL, KHO, BRAH (Brahman), and CIND (Chinese indicine). Blue lines highlight separation time. E) Demographic model for Holstein, Kholmogory and Yakut cattle breeds using GADMA.

We then tried to estimate the time of pairwise divergence between the breeds using SMC++-based coalescent approach. As expected from the historical data, the divergence time of Holstein and Kholmogory was estimated to be around 500 years (Fig. 1A), while the divergence of Yakut lineage was much earlier with a 5,000 years estimation for Yakut-Holstein divergence (Fig. 1C) and 25,000 years for Yakut-Kholmogory divergence (Fig. 1B). The later estimation is likely to be an overestimation (i.e. pre domestication) because the SMC++ does not consider any possible migration events between populations, but estimation of an early Yakut cattle divergence from the common ancestor of Holstein and Kholmogory is in line with an early separation of the Turano-Mongolian cattle from the European taurine breeds.

The second approach, based on the joint allele frequency spectrum (GADMA) returned a quite realistic demographic population model of Yakut, Kholmogory, and Holstein divergence (Fig. 1E). The estimated divergence between Kholmogory and Holstein was around 300 years ago in line with historical data, while the divergence of Yakut cattle from the common ancestor of Holstein and Kholmogory was estimated to be around 4,900 years ago. The best Log-likelihood model also indicated sharp decline of Yakut cattle population around 300 years ago and migrational events from Holstein, Kholmogory and their common ancestor to the Yakut lineage (Fig. 1E). Thus, the very different methods with contrasting assumptions (allowed and not migrational events) point out to a very old (several thousands of years) time of divergence of the Yakut breed from the common ancestor of modern taurine breeds, which was accompanied also by a historically very recent bottleneck of Yakut cattle population. The reality of admixture between European taurines and Yakut cattle is not very clear because we did not observe significant amount of European taurine ancestry in the Yakut genetic pool with ADMIXTURE algorithm (11) but it is also impossible to completely exclude such events.

### Yakut cattle: indicine introgression or ancestral taurine genetic variants

As it follows from our demography and previous studies (11, 20, 62) the Kholmogory and Yakut cattle have different histories, with the Kholmogory being a ‘pure’ taurine breed formed in European Russia in the XVII century as the result of selection applied to admixtured population of imported European cattle and native populations (11). This is confirmed by similar *N_e_* trajectories of the Kholmogory cattle with Holstein (Fig. 1E). The Yakut cattle were likely formed in Asia and demonstrated signatures of possible introgression from indicine cattle (11, 20). To identify genome intervals subjected to potential introgression into the Yakut cattle, a combination of several approaches has been used in this study. We used ‘pure’ taurine (Holstein, Kholmogory) and indicine (African, Chinese, and Indian) breeds as reference populations in the RFMix analysis, which suggested that the Yakut cattle contains ~86.5% of the ‘taurine-like’ and ~13.5% of the ‘indicine-like’ genome. On the other hand, the TreeMix analysis, using a total of 18 cattle breeds and five bovinae species did not detect any introgression between the Yakut cattle and other populations used (Fig. 2). On the TreeMix tree the Yakut cattle was placed close to the Hanwoo and Yanbian breeds and the Kholmogory close to Holstein, which is consistent with previous studies (11, 20). We observed an introgression from banteng to Chinese indicine cattle breeds consistent with the previous report (28) as well as multiple additional introgression events. The *f3* statistics calculated on population triples using the Yakut cattle as a target, various indicine cattle breeds and five related species as source populations did not result in a significant Z-score (Supplementary Information 4a). The *D* statistics (bovinae species, indicine, Yakut, Yak) results, however, suggested admixture between the Yakut and indicine cattle as well as with taurine breeds (Supplementary Information 4b, Supplementary Fig. 1).

**Figure 2.**
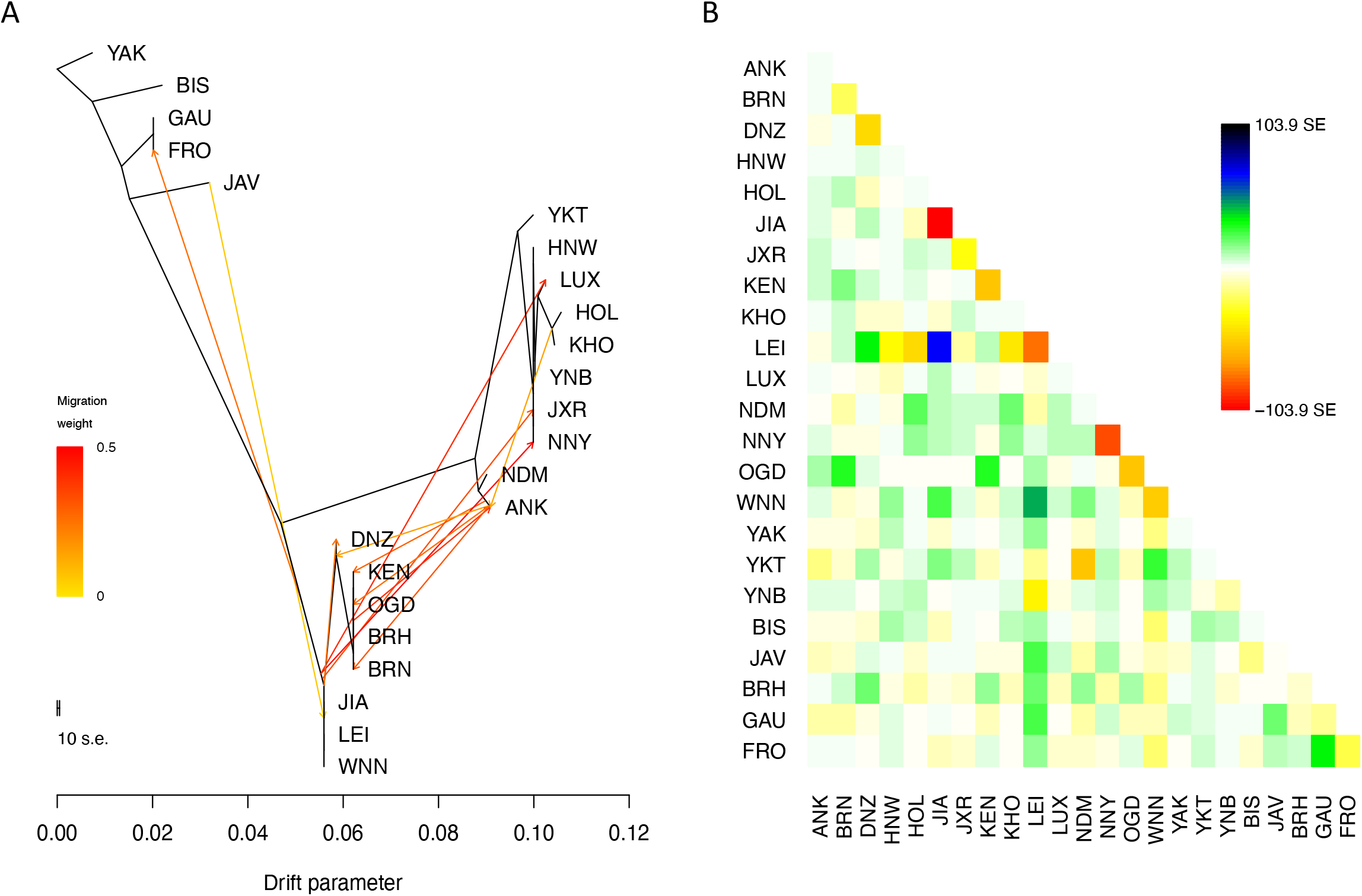
A) The maximum-likelihood tree generated by TreeMix (up to the migration 12). The scale bar shows ten times the average standard error of the entries in the sample covariance matrix. Abbreviations of the populations are as follow: BIS – Bison, GAU – Gaurus, FRO – Gayal, JAV – Banteng, YKT - Yakut, HNW – Hanwoo, LUX – Luxi, HOL – Holstein, KHO – Kholmogory, YNB – Yanbian, JXR – JaxianRed, NNY – Nanyang, NDM – nDama, ANK – Ankole, DNZ – Dianzhong, KEN – Kenana, OGD – Ogaden, BRH – Brahman, BRN – Boran, JIA - Ji’an, LEI – Leiqiong, WNN – Wannan. B) Residuals of the maximum-likelihood phylogenetic tree. As implemented in TreeMix, the residual covariance between each pair of populations i and j is divided by the average standard error across all pairs. This scaled residual in then plotted in each cell (i,j). Colours are described in the palette on the right. Residuals above zero represent populations that are more closely related to each other in the data than in the best-fit tree and thus are candidates for admixture events (SE – standard error).

To investigate possible reasons for discrepancies between the RFMix and the TreeMix results we extracted four sets of SNPs with high derived allele frequencies in the Yakut, Kholmogory, Holstein, and Hanwoo genomes but with low frequency of the same allele in the other three breeds (one breed MAF>=0.7; other three breeds MAF<=0.1; autosomal). The number of such SNPs was 13,257 in the Yakut cattle but much lower in the other breeds: 758 for Hanwoo, 103 for Kholmogory, and 311 for Holstein. Of the 13,257 Yakut SNPs the majority (9,809 (73.92%)) overlapped the Yakut cattle genome intervals identified as ‘indicine-like’ by the RFMix analysis. To look into the evolutionary history of the high-frequency Yakut alleles we focused on all 48 missense mutations present in the 13,257 SNP set (Table 2; Fig. 3). Of these, 37 Yakut cattle high-frequency derived alleles were absent from the European taurine cattle (Holsten and Kholmogory) but preferably had high frequencies in the indicine cattle breeds. Furthermore, 30/37 derived alleles were found in at least one additional bovinae species suggesting that they could be present in the ancestral taurine genome and could be eliminated from at least two European taurine breed (Holstein and Kholmogory) genomes due to selection or drift. To check this hypothesis we looked into the orthologous positions in proteins of up to 82 evolutionary distinct animals (ranging from mammals to fish) and found that for 10/30 positions the Hereford reference (*Btau6*) amino acids matched the amino acid preferably found in the orthologous position of other animals, while for the 19/30 positions the Yakut cattle amino acid was overrepresented in other animal proteins. These observations coupled with the TreeMix and *f3* statistics results support the hypothesis that most of the high-frequency Yakut cattle SNPs are either absent or with low frequencies in the European taurine cattle could represent alleles eliminated from the European taurine cattle. For the final check we looked for the presence of these 48 missense SNPs in the non-European taurine breeds and found 37/48 Yakut cattle alleles in Hanwoo, 46/48 in the Chinese and 42/48 in the African (nDama) taurine breeds confirming our hypothesis that these variants could be present in the ancestral taurine genome.

**Table 2.** High-frequency derived alleles in Yakut cattle genomes

| SNPs | Statistics |
| --- | --- |
| **High-frequency allele in Yakut cattle genomes\*** | **13,257** |
| Yakut cattle-specific | 263 |
| Low frequency in European taurine genomes** | 4,866 |
| Absent from European taurine genomes | 8,128 |
| Found in indicine genomes | 7,546 |
| Found in bovinæ genomes | 5,760 |
| Found *only* in African, Chinese, and Asian taurine genomes | 2,018 |
| Found *only* in Hanwoo genomes | 184 |
| Found *only* in indicine genomes | 166 |
| **Missense alleles** | **48** |
| Found in Chinese taurine genomes | 46 |
| Found in African taurine genomes | 42 |
| Found in indicine genomes | 39 |
| Found in Hanwoo genomes | 37 |
| Absent from European taurine genomes | 37 |
| Found in bovinæ genomes | 30 |
| Evolutionary conserved (Yakut cattle allele) | 19 |
| Evolutionary conserved (European taurine allele) | 10 |
| Yakut cattle-specific | 1 |

\*Allele frequency $\geq 70\%$ in Yakut cattle genomes and $< 10\%$ in Hanwoo, Holstein and Kholmogory cattle genomes; \*\* $< 10\%$ in Holstein, and Kholmogory cattle genomes.

**Figure 3.**
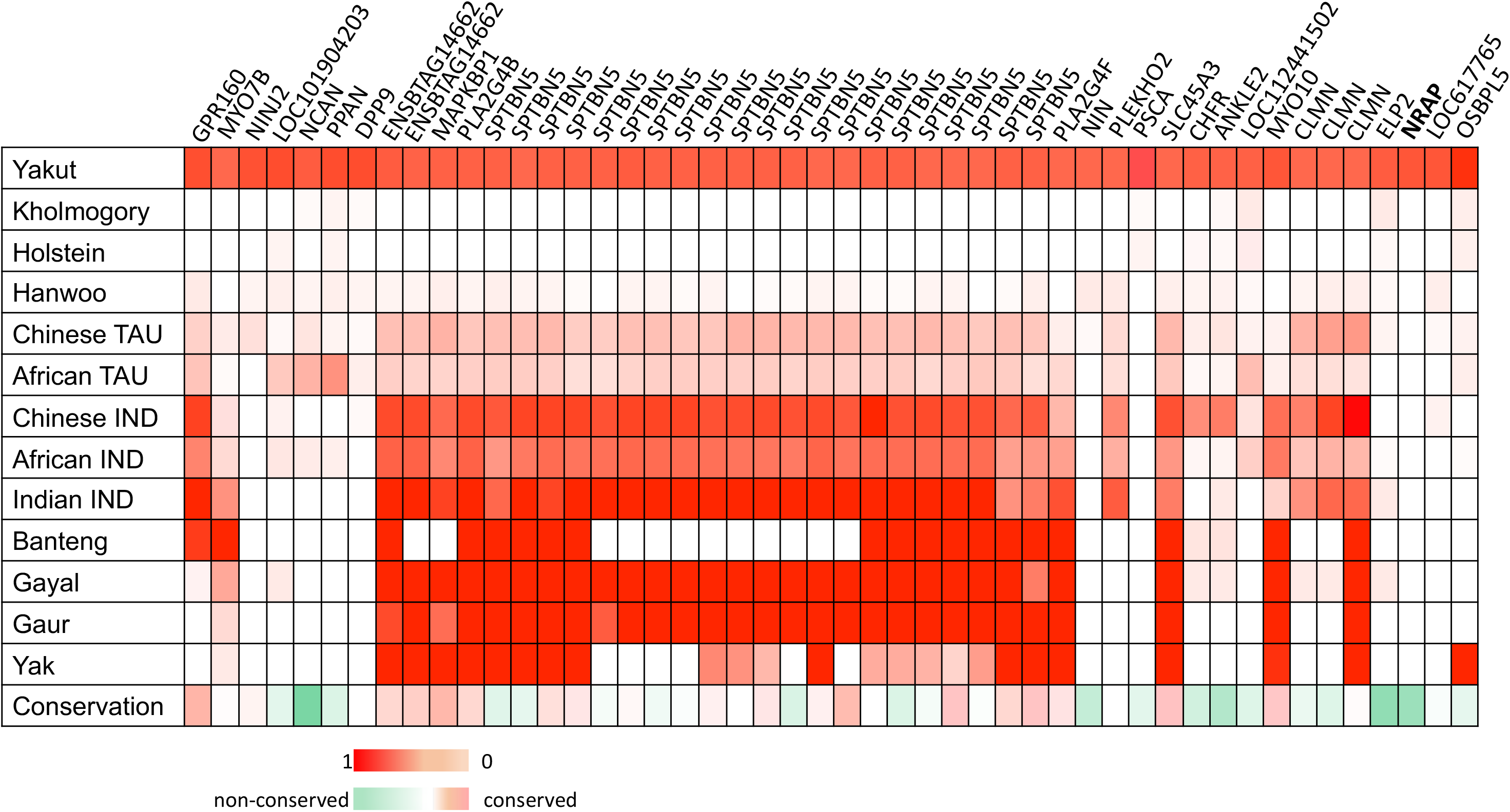
Forty eight missense SNPs with high frequency alleles present in Yakut cattle (>=70%) but with low frequency of the same allele in Holstein, Kholmogory, and Hanwoo (<=10%); shades of red represent allele frequency and conservation score for Yakut cattle allele encoded amino acids more often found in orthologous positions of other mammals, while shades of green indicate those Yakut cattle allele encoded amino acids that are found rarely in other species; TAU – taurine; IND – indicine.

We detected only one Yakut cattle missense derived allele not found in the indicine, taurine cattle or other bovinae species, in the NRAP protein. This H100Q change was highly dissimilar to the orthologous amino acids found in other animals. The fact that all 48 SNPs represent alleles different from the reference genome (Hereford) suggests that they are likely to be absent from both the dairy and beef European taurines. Indeed, we found only 331 SNPs for which the Yakut cattle had a high-frequency allele identical to the reference Hereford genome while Kholmogory, and Holstein had high-frequency derived alleles. None of these 331 SNPs were missense mutations.

### Putative ancestral taurine alleles absent or nearly absent in the European taurine breeds

We used the whole set of 13,257 high frequency Yakut cattle SNPs to identify putative regions of the ancestral taurine genome that are present in high frequency in the Yakut cattle but are absent from the European taurines. Among the 13,257 SNPs we found 263 alleles present in the Yakut cattle only, 4,866 alleles found in a low frequency (<10%) in the European taurines and 8,128 alleles absent from the European taurine set. Of the 8,128 SNPs, 7,546 (92.83%) were found in indicine breeds, however only 166 (2.00%) of them were indicine-specific, while 5,760 (70.87%) were also present in at least one additional bovinae genome, 2,018 (24.82%) additional SNPs were present in either the African or Chinese taurines, and 184 (2.26%) were shared with Hanwoo only (Table 2; Fig. 4 and Supplementary Fig. 2). These results reassured us that the ‘indicine-like’ genome regions of the Yakut cattle to a large extent, if not all, represent ancestral taurine genome regions with alleles mostly eliminated or present in a low frequency in the European taurine genomes. As a final test we identified all intervals in the Yakut cattle genome containing 1.67 million ‘indicine-like’ alleles as assigned by the RFMix analysis, intersected them with bovinae, African, Chinese taurine genomes and Hanwoo, then eliminated all intersected intervals except for those which contained indicine-specific alleles (155,542 SNPs) and repeated the RFMix analysis. It showed that the filtered set had 2.5% of the ‘indicine-like’ genome suggesting that the removed intervals were responsible for the majority of the ‘indicine-like’ part of the Yakut genome.

**Figure 4.**
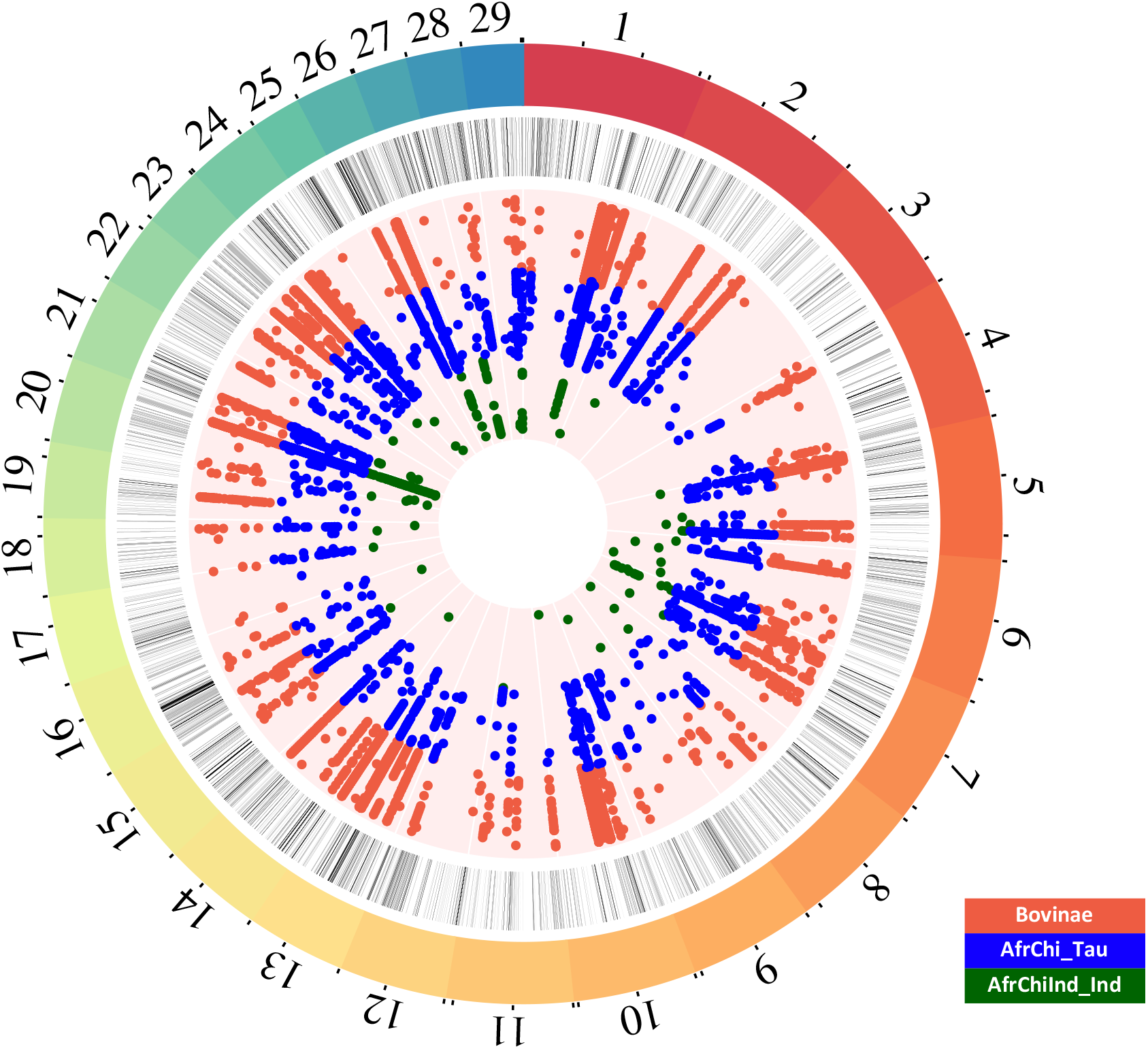
High frequency SNPs in Yakut cattle but low frequency/absence of the same allele in Holstein, Kholmogory, and Hanwoo (total of 13,257 SNP). Each circle from the periphery to the centre shows the following datasets: bovine autosomes, indicine-like genome intervals from the RFMix analysis in different shades of grey (light to dark indicating indicine-like interval frequencies in the Yakut population ranging from 50% to 100%), Yakut alleles found in bovinae species in red, in African and Chinese taurine breeds in blue, and exclusively in Indian, African and Chinese indicine breeds in green.

To understand the role of genes found in the ancestral taurine genome segments, we extracted all genes found in the ancestral and European taurine genome intervals (gene total length overlap >=60% for a gene to be assigned to an either set). A total of 1,639 (6.7%) genes were assigned to the ancestral and 19,205 (78.2%) genes to the European taurine segments. GO enrichment analysis of the genes in ancestral segments revealed the term “*response to stimulus*” (q-value = 0.0) and “*metabolic process*” (q-value = 0.0) among others (Fig. 5; Supplementary Information 5a). Moreover, the genes from the ancestral segments were enriched in the “*immune system process*” pathway (q-value = 0.0; Fig. 5) including the MHC class II antigens. The top DAVID cluster (enrichment score 3.68) included multiple MHC class II antigens, immune genes (e.g. *IL36A* and *IL36B*), heat shock proteins (*HSPA1A* and *HSPA1L*) and the insulin receptor (*INSR*; Supplementary Information 5b). Moreover, we selected four GO categories potentially involved in adaptation to extreme environments: response to stress, response to pain, neuropathy, and actin binding (Supplementary Information 6). The GO category response to pain resulted significantly enriched in genes found in ancestral segments identified by RFMix (l2fold = 1.21, p-value = 5.00e-03, q-value = 1.00e-02; 10,000 permutation test).

**Figure 5.**
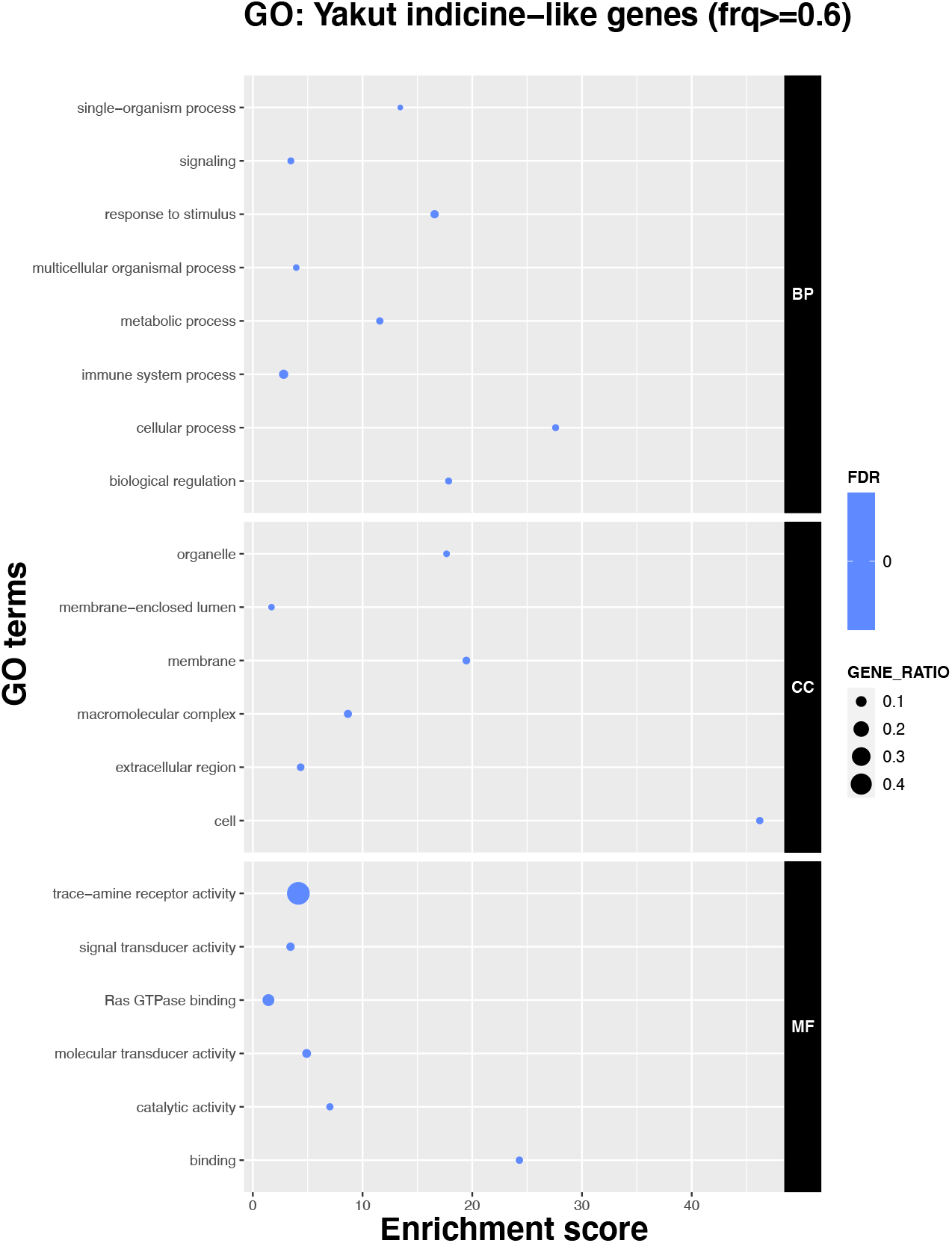
GO analysis of genes overlapping ‘indicine-like’ genome intervals (>=60% of the total gene length) as identified by RFMix.

We then looked closer into the 25 genes that had high-frequency derived missense alleles in the Yakut genomes (Fig 3). A gene with the largest number of such variants was the actin-binding *SPTBN5* gene containing 21 missense mutations. Not surprisingly, the region containing the *SPTBN5* also demonstrated the highest F_*ST*_ value between the Yakut cattle and all taurine breeds from the Run 7 of the 1000 Bull Genome project suggesting that this region is quite different between the taurine breeds and the Yakut cattle. When all breeds of the Run 7 were contrasted against the Yakut cattle *SPTBN5* was still on top of the list (top fourth gene position, with the BOLA-DQB MHC class II antigene being the number one; Supplementary Information 7). We found high-frequency derived alleles in other actin-binding genes, including the *MYO10, CLMN*, and the *NRAP*. Among these the *MYO10* also contained a unique missense mutation in the Yakut horse (63), suggesting its probable contribution to the local adaptation of distinct domesticated species to harsh Siberian climates.

### Yakut-specific NRAP missense mutation

The *NRAP* variant was the only Yakut-cattle specific missense mutation which was not observed in other cattle breeds nor the bovinae species in our set. Out of 19 Yakut cattle individuals with high coverage in this region (no. reads >5) 12 were homozygous for the derived allele while seven were heterozygous. We checked this position in the complete 1000 Bull Genome Run 7 dataset (3,767 individuals, indicine and taurine breeds). The frequency of the reference (G) allele was 0.995089 while the frequency of the derived (T) allele was 0.00491107. The only four additional (to our Yakut cattle samples) animals which contained the same derived allele were four Yakut cattle samples obtained from the NCBI SRA achieve, suggesting that this genetic variant is Yakut cattle-specific.

The nebulin-related-anchoring protein (NRAP) is a highly evolutionary conserved actin-binding cytoskeletal protein expressed exclusively in the skeletal and cardiac muscles, and is involved in myofibrillar assembly and force transmission in the heart (64). Our phylogenetic comparison shows that the H100Q variant observed in the Yakut cattle is rare in other mammals (found in 16/113 species), but is preferably present in species that either hibernate, e.g. little brown bat and other hibernating bats and mouse lemur or enter seasonal torpor e.g., cape golden mole and tree shrew. Our statistical analysis using phylogenetic logistic regression (65) considering phylogeny of the 113 species from nine mammalian orders (66) shows phylogeny-independent significant correlation (R^2^ 0.77, p-value = 2.72e-06, alpha 0.00011) between the presence of the glutamine (Gln, Q) variant and cold adaptation, hibernation or ability to enter torpor (Fig. 6, Table 3). The four species that don’t hibernate or enter torpor but still have the glutamine variant are the walrus, sea lion and two seals. Interestingly seals and sea lions are capable of slowing heart rate to several beat per minute while diving (67, 68) while walrus is cold adapted suggesting that the same adaptation as in the Yakut cattle could be found in other large animals.

**Figure 6.**
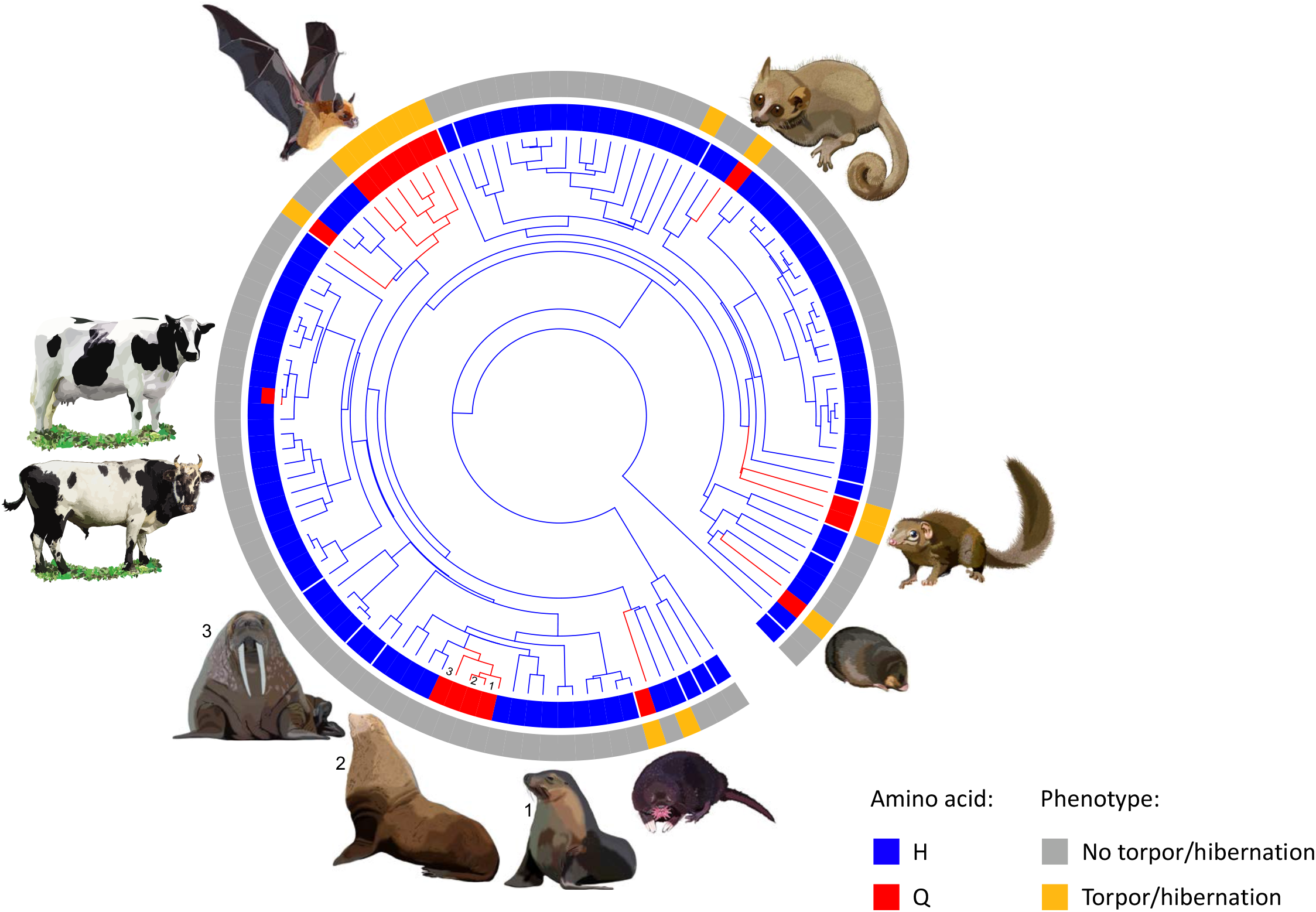
Phylogenetic tree of 114 mammals from nine phylogenetic orders (100). Carriers of the common histidine (H) amino acid are in blue while the carries of the glutamine (Q) variant are in red (cattle has both blue and red indicating the Yakut cattle specific H100Q mutation); the outer circle in orange highlights species that either hibernate or enter torpor (Supplementary Information 17). Images are representative of the species that have the glutamine variant, Yakut cattle (glutamine/histidine) and Kholmogory cattle (histidine).

**Table 3.** Association between allelic variants at NRAP residue 100 and habitat across 113 mammalian species.

| Residue | Amino acid | No. species |  | R<sup>2</sup> | P value* |
| --- | --- | --- | --- | --- | --- |
|  |  | Torpor/hibernation | No torpor/hibernation |  |  |
| 100 | His | 2** | 95 | 0.77 | 2.72 x 10<sup>-06</sup> |
|  | Gln | 12 | 4 |  |  |

\* phylogenetic logistic regression test. \*\*tenrec – hibernates when it is hot rather than cold and the alpine squirrel. One species, the golden hamster, has not been included in the analysis as the only hibernating species which shows arginine (Arg), a descendant of glutamine (Gln) at the NRAP 100 position.

There are multiple independent lines of evidence suggesting that the highly evolutionarily conserved NRAP protein could contribute to climate adaptations in various animal groups. For instance, in the Australian bearded dragon *NRAP* is differentially expressed in the skeletal muscle and heart before and during hibernation (69) while in the alligator *NRAP* expression was downregulated in the heart in response to hypoxia (70). In the glass eel fixation of an alternative *NRAP* allele was associated with latitude and river water temperature in a landscape genomics study (71). Interestingly, in a recent environmental GWAS study (72) variants in the *NRAP* gene were associated with environmental adaptations in the US Simmental cattle.

Human and mice studies provide a key to likely mechanism of the *NRAP* contribution to cold climate adaptations. In mice *NRAP* is downregulated during dilated cardiomyopathy (DCM) (73); human patients with DCM express homozygous *NRAP* mutations (64). During DCM, pumping ability of the left ventricle of the heart increases leading to issues with pumping blood out of the heart (74) suggesting that the heart force transmission function of *NRAP* in the Yakut cattle could lead to more efficient blood pumping during the winter period therefore facilitating energy saving similar to species that either lower metabolic rates during the winter period, enter torpor or hibernate (Fig. 6 and Fig. 7).

**Figure 7.**
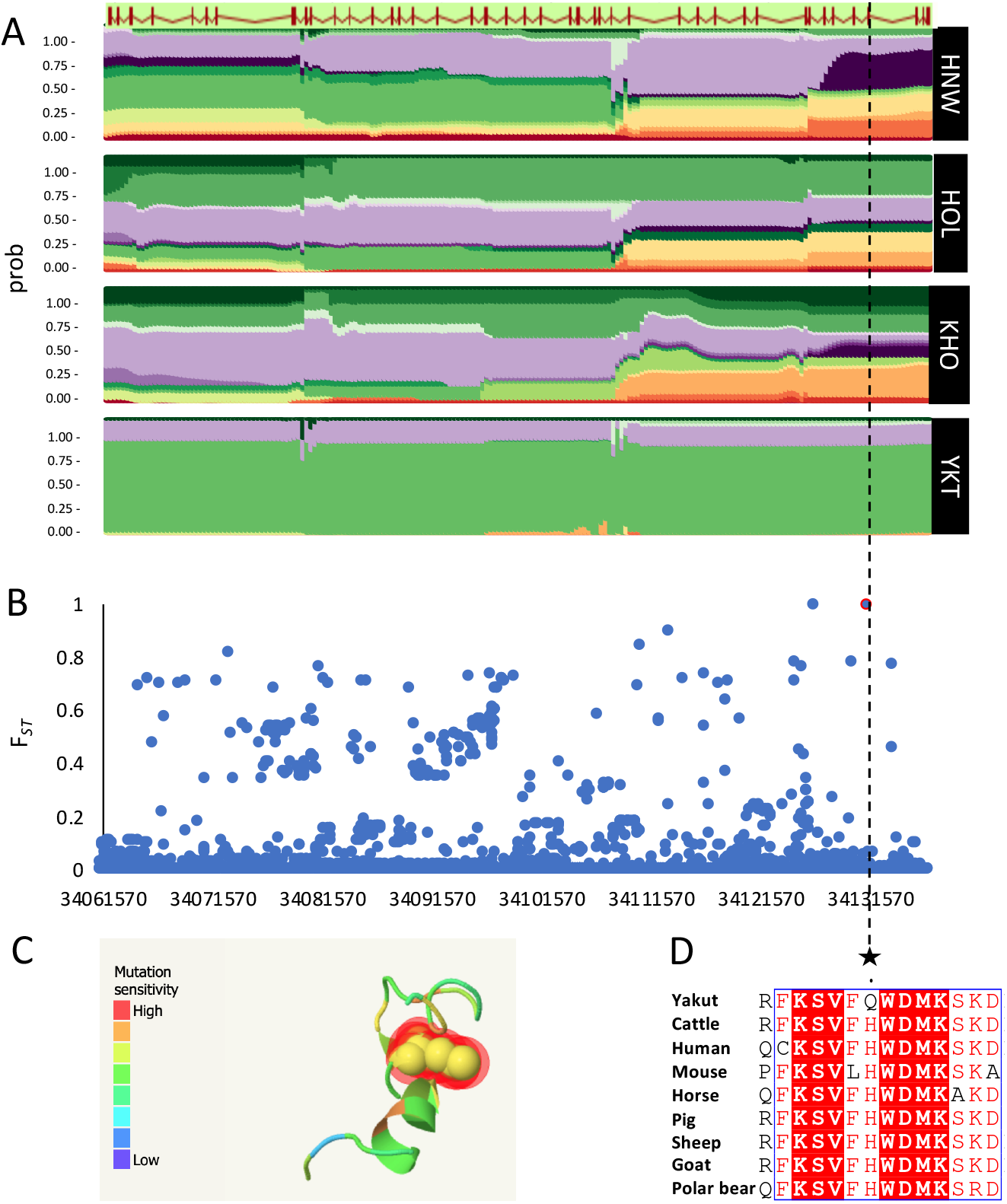
SNPs in the *NRAP* gene. A) Haplotype diversity in the *NRAP* gene for Hanwoo (HNW), Holstein (HOL), Kholmogory (KHO), and Yakut (YKT) breeds. B) F*_ST_* analysis of SNPs from the 1000 Bull Genomes Run 7 in the Yakut population against all other individuals; C) Sensitivity of the H100Q change modelled in the NRAP protein; D) Extract of local alignment of the exon 4 of the *NRAP* gene showing the H100Q mutation and adjacent amino acids in the Yakut cattle (Yakut), reference Hereford genome (Cattle) and seven mammals.

### Copy number variant detection in the Yakut, Kholmogory, Holstein, and Hanwoo breeds

Copy number variants (CNVs) together with SNPs are a major source of genomic variation contributing to evolution and adaptation (75). To identify CNVs and CNV regions (CNVRs) which could co-evolve with adaptation of the Russian cattle breeds to harsh climate conditions, we utilized sequences of four breeds (Yakut, Kholmogory, Holstein, and Hanwoo). A total of 60,338 autosomal CNVs were detected in the four-breed set, which were then merged into 17,736 CNVRs. The largest shared fraction of Kholmogory CNVRs was with Holstein (2.48 Mbp) and Yakut CNVRs with Hanwoo (2.51 Mbp), confirming known breed relations (Fig. 8A). A total of 588 CNVRs (total length: 2.42 Mbp) were breed-specific (had <10% length overlap with the CNVRs from other three breeds) for the Yakut, 743 (3.73 Mbp) for the Kholmogory, 1,408 (5.42 Mbp) for the Hanwoo and 2,464 (8.97 Mbp) for Holstein cattle (Supplementary Information 8a, Fig. 8a). To check if the unique CNVRs in the Russian (especially Yakut) cattle could result from ancestral taurine genome, we additionally identified 9,960 CNVRs in 26 samples of pure indicine (African, Chinese, and Indian) breeds. Overall, the Yakut and Kholmogory cattle had the same fraction of their unique CNVR lengths overlapping the indicine CNVRs (34%), while Hanwoo had the largest fraction (51%) and Holstein had the lowest (23%; Supplementary Information 8a).

**Figure 8.**
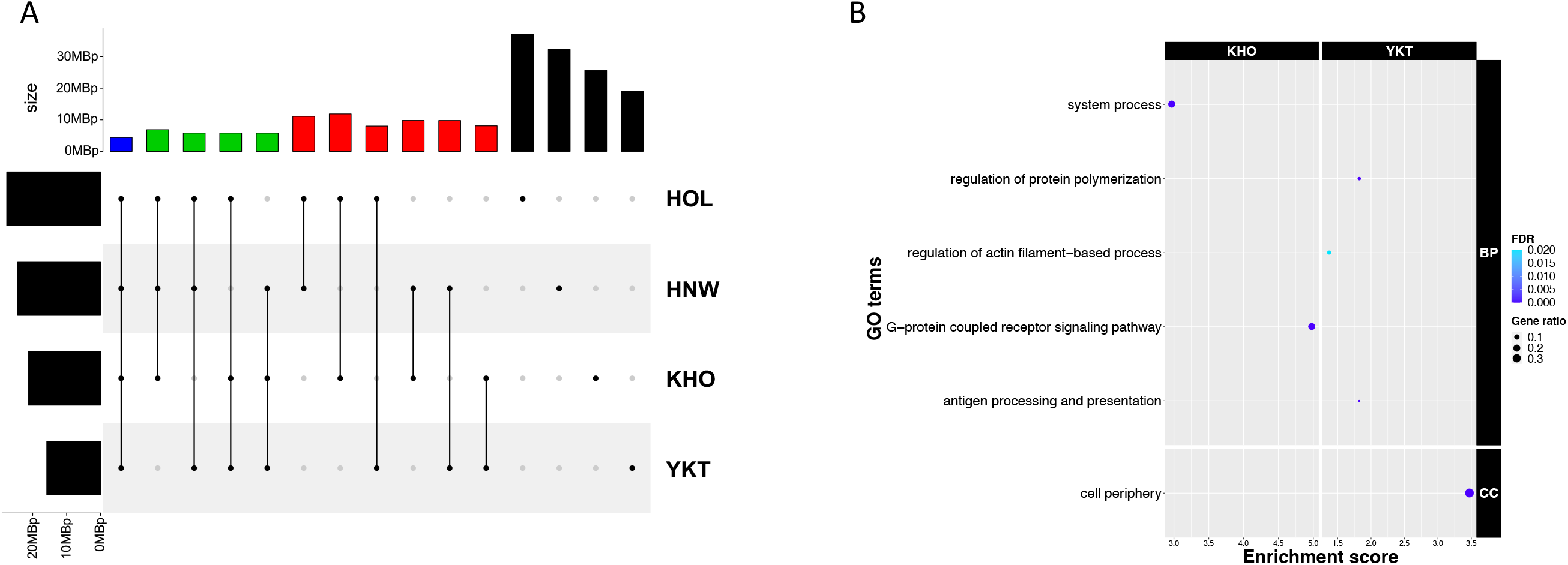
Summary of copy number variant regions (CNVRs) in Yakut, Kholmogory, Holstein, and Hanwoo breeds. A) intersection plot of CNVRs for the four breeds (HOL – Holstein, HNW – Hanwoo, KHO – Kholmogory, YKT – Yakut); colours indicate: black, breed-specific CNVRs; red, intersection between two breeds; green intersection among three breeds; blue intersection among all four breeds. B) GO analysis of genes in breed-specific CNVRs in Yakut (YKT) and Kholmogory (KHO) (FDR<0.05); only unique GO terms present in either Yakut or Kholmogory breeds were reported.

**Figure 9.**
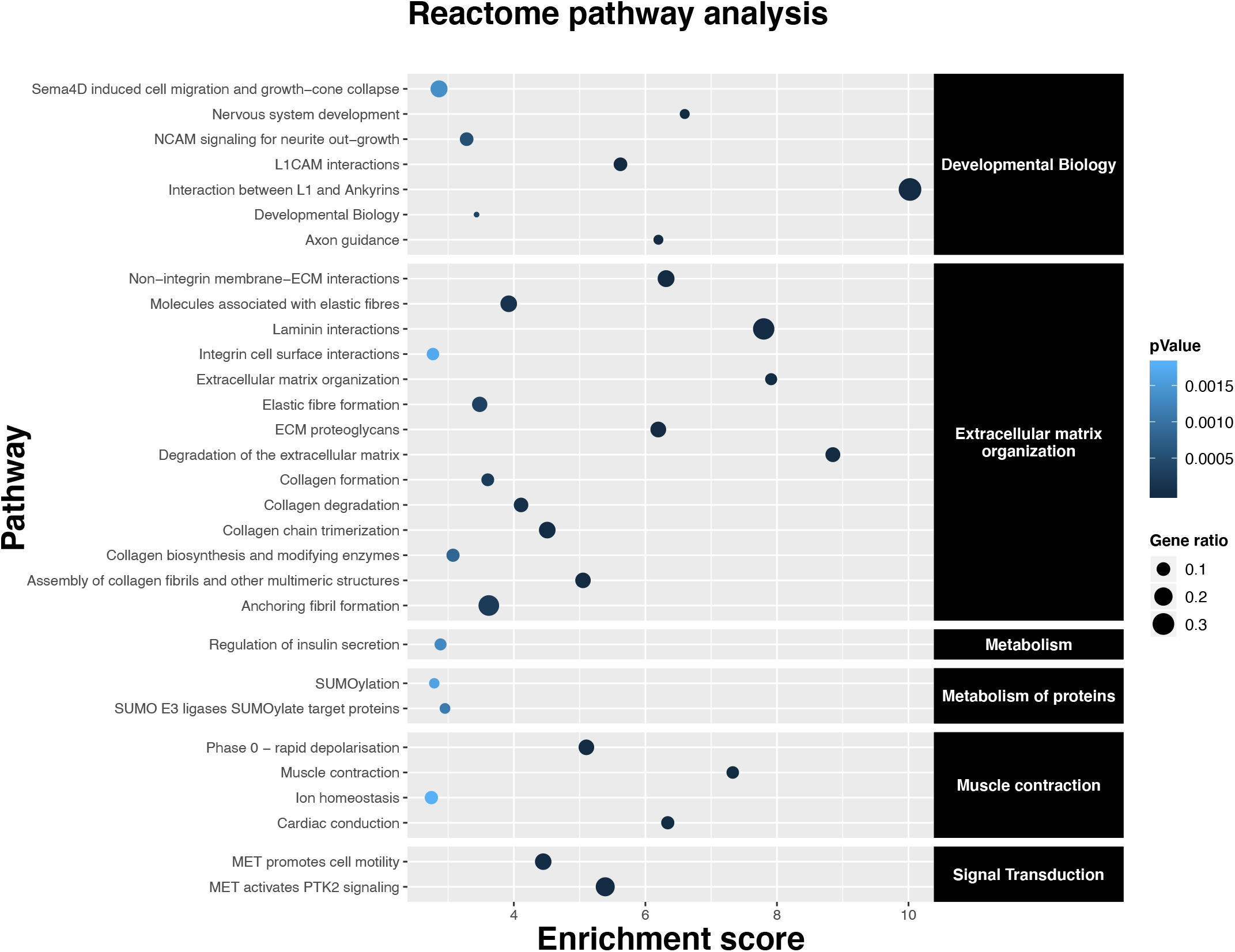
Reactome pathway enrichment analysis for Yakut cattle genes identified to be under selection by the dN/dS analysis (FDR<0.05).

To reveal possible contribution of CNVRs to the breed-specific biology and adaptations we looked into 1,887 genes found in the four breed-specific CNVRs of which 1,261, 279, 208, and 139 genes were found in the Holstein, Hanwoo, Kholmogory, and Yakut CNVRs, respectively (Supplementary Information 9a). Gene Ontology (GO) enrichment analysis highlighted distinct pathways being enriched in these gene sets, with the largest number (25) found in the Holstein cattle (Supplementary Information 9a). The Yakut cattle had the second largest number of GO categories (four) of which *regulation of actin filament-based processes*, and *antigen processing and presentation* could potentially contribute to adaptation to local environments. The Kholmogory showed enrichment in two GO categories, *G-protein coupled receptor signalling pathway* and *system process*, while Hanwoo had only one GO category enrichment, the *ion binding*. There were 299 CNVRs shared between the Yakut and Kholmogory breeds, of which the CNVR covering >90% of *ACTG1* (actin gamma 1) gene was the most frequent (15% and 11% in Kholmogory and Yakut, respectively). Another actin gene, *ACTA1* (alpha-skeletal actin) was initially reported in the Yakut-specific CNVR, with one Kholmogory individual having CNV in the same region. A manual check of BAM files revealed that more Kholmogory samples followed the same increased coverage/heterozygosity pattern, suggesting that this CNV is more widely spread in these two breeds, while none of the Holstein and Hanwoo samples showed this pattern and had actin genes in their CNVRs. The gene network analysis (Supplementary Fig. 3) shows that *NRAP* affected by the Yakut-specific missense mutation (see above) directly interacts with *ACTA1* gene supporting assembly of actin filaments in both skeletal muscles and the heart. Additional Yakut cattle-specific CNVRs involved multiple olfactory receptor genes, *CYP4A11* (cytochrome P-450 4A11) involved in lipogenesis and growth traits. This CNVR has been previously reported in various Chinese native taurine cattle breeds (Jaxian, Quinchuan, Nanyang, Jinnan, Luxi, and Chinese Red Steppe (76)).

The Kholmogory cattle unique CNVRs covered 54 annotated genes among which 17 were olfactory receptors, *DEFB1* (defensin beta 1) antimicrobial peptide, *IFITM2* (interferon induced transmembrane protein) both involved in the bovine innate immunity, *PLA2G2D3* (phospholipase) involved in inflammation and immune response, *C8G* (complement C8 gamma chain) belonging to the lipocalin family and participating in the formation of the membrane attack complex, *MYLPF* (myosin light chain) associated with meat tenderness and intramuscular fat (77), *SHOX* (short stature homeobox) regulating elements of growth and development and its mutations are associated with skeletal atavism (78).

Among the genes in the breed-specific CNVRs overlapping indicine CNVRs (356 (1.45%)) 112, 109, 81, and 54 were present in the Holstein, Hanwoo, Kholmogory, and Yakut breeds, respectively (Supplementary Information 8b). GO enrichment analysis highlighted the term “*detection of stimulus*” in both Yakut and Kholmogory breeds, and the term “*MHC protein complex*” in the Yakut cattle (Supplementary Information 10a).

### Genomic regions under selection in world northernmost cattle breeds

In an attempt to decipher genomic regions under selection in Russian cattle breeds and thus better understand their unique adaptations, we utilized two independent approaches: one focusing on the identification of haplotypes subjected to selection in a population (HapFLK) and another looking for the ratio of non-synonymous to synonymous substitutions within a gene (dN/dS) while considering the background genome mutation rate to differentiate between the signatures of positive and negative selection. In the latter approach, we first analyzed all SNPs in each breed (MAF =<0.1) and then reanalyzed the dataset focusing on SNPs present in more than half chromosomes in each population (MAF>=0.6). In fact, we were interested in finding genes under selection with a likely effect on the whole population rather looking at all variation within a breed. Indeed, most genes found to be under selection when considering a more stringent allele frequency were also reported under selection when more SNPs were considered (Supplementary Information 11). In addition, we used information about 13,257 high-frequency SNPs with derived alleles in Yakut cattle to support our findings together with the top 1% fixation index of genetic differentiation (F*_ST_*) intervals in Yakut and Kholmogory breeds calculated against the *pure taurine*, *all taurine* and *all breeds* from the 1000 Bull Genome Project. Calculation of functional category enrichments within these sets were independently estimated using the GO categories, DAVID functional clusters, and the Reactome. Overall, we identified 282 regions under selection in the Yakut and 219 in the Kholmogory cattle breeds using the HapFLK analyses (q-value <0.01; Supplementary Information 12), four and three genes under positive and 247 and 62 genes under negative selection using the dN/dS in the Yakut and Kholmogory breeds, respectively when applying the 60% MAF filter (Supplementary Information 11b). There were 14 and four genes in the Yakut and Kholmogory cattle, respectively from the HapFLK selected regions which were also picked up by the dN/dS analysis. Of these, genes *SPTBN5* was also found in the top 1% F*_ST_* intervals in the Yakut cattle, while *HID1* (involved in regulation of neuropeptide sorting) in the Kholmogory cattle.

As previously shown, adaptation to harsh environments requires proper responses to external stimuli (79). Both the Yakut and Kholmogory breeds demonstrated significant enrichment for the *response to stimulus* GO category in the HapFLK regions, supported further by the dN/dS analysis of the Yakut cattle genes. Interestingly, in the Yakut cattle this GO term was significantly overrepresented in genes containing high-frequency SNPs absent from the European taurines, suggesting that the genetic profile of selection between the two breeds is different. One of the strongest HapFLK signals in the Yakut cattle was observed for the gene *ZNF622*, which was recently established as an antiviral protein in humans (80). Glycophorins (*GYPB* and *GYPA*), which determine MN and Ss blood types in humans (81) and define resistance to parasites, including malaria (82), demonstrated high F_*ST*_ between the Yakut cattle and the pure taurine cattle with the *GYPB* being on top of the F_*ST*_ gene list (Supplementary Information 7) and the positive selection list in Yakut cattle from the dN/dS analysis (MAF =<0.1; Supplementary Information 11a). Our finding of a convergent *NRAP* amino-acid change in the Yakut cattle shared with multiple cold-adapted/hibernating species suggesting that the Yakut cattle could be capable of slowing metabolism during the coldest winter periods, was supported by the GO *metabolic process* (q-value = 0.00; Supplementary Information 13a) and *integration of energy metabolism* (q-value= 0.04; Supplementary Information 13c) Reactome terms enrichment in the HapFLK Yakut gene set. This adaptation would require changes to the heart’s ability to distribute blood efficiently despite cold environmental temperatures. Indeed, the *cardiac conduction* Reactome pathway was overrepresented (q-value = 4.94E-05; Supplementary Information 14a and Fig. 9) in the dN/dS gene set supported by the DAVID overrepresented cluster (enrichment score 2.00; Supplementary Information 15b) combining terms *cardiac muscle contraction, dilated cardiomyopathy, hypertrophic cardiomyopathy*, etc. when the Yakut cattle F_*ST*_ was calculated against the pure taurine breeds from the 1000 Bull Genome Project dataset and the same cluster with enrichment score of 1.32 when compared to all breeds (Supplementary Information 15b). These categories included a common gene *RYR2*, which is central to myocardial excitation contraction coupling (83).

An important organ contributing to thermogenesis is the brown adipose tissue (84, 85). In the Yakut cattle the third strongest HapFLK signature of selection was in the area of the pleiotrophin (*PTN*; q-value = 4.42E-10; Supplementary Information 12), a key gene in preserving insulin sensitivity, driving adipose tissue lipid turnover and the regulator of energy metabolism and thermogenesis (86). In line with this observation we found that the *regulation of insulin secretion* term was enriched (q-value = 0.04; Supplementary Information 14c) in the Yakut dN/dS gene set (Fig. 9). This was further supported by the HapFLK analysis highlighting the Reactome categories *regulation of insulin secretion* (q-value = 0.01) and *integration of energy metabolism* (q-value = 0.04; Supplementary Information 13c). Intriguingly, we identified two out of three candidate genes for cold/diet adaptation in native Siberian human populations (*PLA2G2A* and *ANGPTL8*) (87) involved in lipid metabolism in the Yakut cattle HapFLK set. Another mechanism of reaction to extreme cold temperatures in mammalian cells is the cytoskeletal stabilization and/or disassembly including microtubules and actin filaments. In hibernating mammals this is required for a rapid cytoskeleton reorganization during return from torpor to euthermy (88). We found multiple cytoskeleton proteins in selected intervals of the Yakut cattle genome, including highly-enriched DAVID clusters of *spectrin/alpha-actinin*, etc. (enrichment score 6.22); *myosin, actin-binding, microfilament motor activity, actin filament binding*, etc. (enrichment score 5.14) in the dN/dS set (Supplementary Information 14b). Genes with the highest selection signals included actin-binding *SPTBN5* and *MYO10*, both of which were highlighted by our high-frequency SNP analysis. The *SPTBN5* is among 12 genes under selection in multiple Arctic and Antarctic species (18), while *MYO10* contained Yakut horse specific mutations (63). Interestingly, the actin-binding genes were also enriched in selected intervals of Kholmogory and Hanwoo breeds, but at much lower enrichment rates (enrichment score 2.95 in Kholmogory and 1.92 in Hanwoo) and the gene sets were different to Yakut cattle (Supplementary Information 14b). Another related group of cytoskeleton-related proteins under selection in the Yakut cattle were *pleckstrin homology-like domain* proteins in the dN/dS (enrichment score 2.80; Supplementary Information 14b) and HapFLK (enrichment score 1.63; Supplementary Information 13b) analyses. These domains are found in proteins that link the cytoskeleton to the cell membrane (89). One of these genes, *FARP1* shows the highest HapFLK signal in the Yakut cattle (q-value = 0.0; Supplementary Information 12). An important organ of thermoregulation is the brain. It was shown (90) that the extracellular component proteins play role in axonal guidance, synaptogenesis (collagens) and were overexpressed in the cerebral cortex during deep hibernation of ground squirrel. A similar pattern was observed in hypothalamic transcriptome of Djungarian hamster during torpor (91). The Reactome analysis of the dN/dS gene set highlighted multiple extracellular matrix terms enriched in the Yakut cattle, including *elastic fibre formation* (enrichment 3.48; q-value = 0.01); *laminin interactions* (enrichment 7.79; q-value = 3.2E-06), *assembly of collagen fibrils and other multimeric structures* (enrichment 5.05; q-value = 5.16E-04), *collagen formation* (enrichment 3.60; q-value = 0.01), etc. (Fig. 9; Supplementary Information 14c).

#### Gene enrichment for Yakut SNP set absent or in low frequency in the European taurine cattle

To understand probable contribution of the alleles, absent from European taurines, we looked for GO enrichment in the 13,257 SNPs with high frequency in the Yakut cattle. The *actin cytoskeleton* (q-value = 0.0) category was overrepresented in the set of SNPs that were found at low frequency (<=10%) in the European taurines, while the *response to stimulus* and *metabolic process* categories were enriched (q-value = 0.04) in the set of SNPs absent in the European taurine breeds (Supplementary Information 16a). We observed that SNPs absent in the European taurines but found in bovinae species were enriched in the GO category “*phospholipase A2 activity*” (q-value = 0.0), the Reactome pathway *Acyl chain remodelling of PG* (enrichment score = 3.7; q-value = 0.05; Supplementary Information 16c), and “*MHC class I peptide loading complex*” (q-value = 0.0). Interestingly the genes containing Yakut-specific SNPs were enriched in *biological regulation* (q-value = 0.0; Supplementary Information 16a).

## Discussion

Native cattle breeds represent an important cultural heritage, often sharing common history with local human populations. Local breeds could also contain genetic variations useful to properly respond to agricultural needs in light of ongoing climatic changes. They could help understand evolutionary processes that occur in response to human led selection and adaptation of cattle to different (sometimes drastic) environmental conditions. In this study we looked into the genetic history of two unique world northernmost native cattle breeds from Russia: the Kholmogory, and the Yakut cattle. We revealed the molecular layout for distinct genetic profiles of European and Asian taurine cattle lineages, spotted a recent functional convergent evolutionary change in the Yakut cattle genome in response to exposure to climates of Siberia, found candidate genes and pathways which could contribute to cattle adaptations to cold climates, and offered a set of SNP markers for the industry to be tested for genomic selection of cattle in the Northern countries.

Comparison of demographic histories of the Yakut, Kholmogory, and Holstein breeds suggests an early separation of the Yakut cattle from European taurines (represented by Kholmogory and Holstein). The estimated separation time (4,900 years ago) agrees with Payne and Hodges (92) who estimated that domesticated cattle appeared 4,000-5,000 years ago in the Northern East Asia, Northern China, Korea and Japan (92). As this is a post-domestication event with the domestication itself dated to ~8,500-10,000 years ago (1, 2), our data supports a single taurine cattle domestication leading to both the European and Asian taurines. Our results, however, point to a much smaller *Ne* size of the ancestral Asian taurine population compared to a much larger *Ne* of the population leading to European taurine breeds. This could be due to known historical admixture of European cattle population with wild aurochs (93) until the European cattle’s *N_e_* reduced ~2,500 years ago and possibly could account for phenotypic differences between the extant Turano-Mongolian cattle and European taurines. Interestingly, there is a slight gradual increase of the *N_e_* of the Yakut cattle lineage until its drastic decrease ~300 YA, which approximately matches the period of the Yakutian history when in 1620s they came in contact with the expanding Tsardom of Muscovy. The imposed fur tax led to several suppressed Yakutian rebellions between 1634 and 1642, but more importantly, there is evidence that the Yakutian population went down by ~70% between 1642 and 1682 due to smallpox and other introduced infectious diseases (94). Our data suggest that the Yakut cattle population could have gone through a severe bottleneck at nearly the same time corroborating the event in the human history.

A high fraction of SNPs with high frequency in the Yakut cattle, absent from the European taurine breeds, but present in the indicine cattle, could imply historical introgression of these alleles into the Yakut cattle. However, the fact that vast majority of these alleles were also present in different bovinae species means that these variants could belong to the pool of ancestral *Bos primigenius* alleles shared with other *Bos* species. Indeed, we identified a large fraction of alleles with high frequency in the Yakut cattle, but with low frequency in European taurines and Hanwoo, suggesting that these alleles were segregating in the ancestral domesticated population more than 5,000 years ago. The fact that the Yakut and Kholmogory (a pure taurine) cattle had nearly the same fraction of CNVRs overlapping CNVRs from indicine breeds provides another line of evidence against significant introgression of Yakut and indicine cattle. Another possible scenario which could potentially explain high genetic diversity of Yakut cattle at these positions is an ancient introgression with the East Asian aurochs (*Bos primigenius*) that lived in the East Asian region during the arrival of taurine cattle together with several other bovinae species, including yak, banteng, and gaur (3, 4, 28). Kantanen et al. (95) have indeed suggested the near-eastern origins of the ancestral population of the Yakut cattle breed by studies on mtDNA and Y-chromosomal diversity. This scenario, however, cannot explain large fraction of SNPs with a high frequency in the Yakut cattle, indicine cattle, and African taurines, and with a low frequency in European taurines. The fact that nearly two-thirds of missense alleles absent from the European taurines are more similar to orthologous amino acids found in majority of other phylogenetically distinct animal species also implies that these variants could represent the ancestral state. We hypothesise that these putatively ancestral variants could be a rich reservoir of genetic variation important for adaptation of Yakut to extreme cold environment. This is indeed supported by their enrichment in genes from *response to stimulus/pain* and *immune system* categories potentially involved in adaptation to harsh environment. The fact that the *response to stimulus* category is under selection in another Northern Russian breed, the Kholmogory, confirms similar mechanisms of adaptation to cold climates in cattle breeds. Interestingly, the GO category response to stimulus was found to be enriched in a recent whole genome resequencing study of the Finnish and Yakut cattle breeds in the list of genes with large number of missense mutations (96). A large fraction of genes in selected regions in the Yakut cattle genome belong to the actin-binding category; the same category was found enriched in the selected regions of the Kholmogory cattle, suggesting that these genes could contribute to cold adaptation as it was shown previously in (38, 88, 97). This is supported by the fact that the *ACTA1* (alpha actin 1) was the most frequent Yakut CNVR and also present in Kholmogory cattle, while this CNVR was absent from all Holstein and Hanwoo samples. Overall, the CNVRs found in the Yakut cattle were significantly enriched for actin-binding genes, suggesting that this group of genes has been evolved in the Yakut cattle. Another actin-binding gene *SPTBN5*, which had the largest number of high-frequency missense mutations in Yakut cattle (21) compared to other taurine breeds, was shown to be under positive selection in at least three Arctic and Antarctic mammals including the woolly mammoth (18, 98) and in Chinese indicine cattle breeds (27); *MYO10* gene which has unique missense mutation in the Yakut horse (63), suggesting its probable contribution to local adaptation of distinct domesticated species to harsh Siberian climates. Our findings are confirmed by the work of Weldenegodguad et al. (96) who identified genes with >5 missense mutations in the Yakut cattle genome from five samples, and both the *SPTBN5* and *MYO10* were reported in their list; but the limited number of animals in their study was not sufficient to estimate signatures of selection and frequencies of derived alleles in the Yakut cattle genome.

While the actin genes themselves and genes interacting with actin could contribute to multiple biological processes related to adaptation to cold climates (97) the only high-frequency Yakut-cattle specific missense mutation found in the actin-binding gene *NRAP* provides a clue for the likely mechanism. The fact that this H100Q variant, found in every single resequenced Yakut cattle animal was absent from all ~3,800 cattle animals from the 1000 Bull Genome Project, including ancient aurochs samples, suggests its Yakut cattle origin. Moreover, its absence from all bovinae species used in our work as well, suggests its relatively recent appearance in the Yakut cattle dating back to maximum 5,000 years ago, after separation from the European taurine branch. However, as the Yakut people have migrated to Yakutia in the 13^th^ century and other Asian, including the Turano-Mongolian breeds (Hanwoo), lack this variant as well it could form as late as ~800 years ago in Yakutia followed a very fast expansion in the Yakut cattle. While absent in cattle breeds and phylogenetically close bovinae species, the exact same amino acid change has formed independently at least six times in mammalian evolution, in species lowering their metabolism in response to cold, including hibernating bats and lemurs, cold adapted species, like walrus and sea lions. Interestingly, a potential contribution of a highly evolutionary conserved NRAP protein (99) to climate adaptation is supported by findings in a range of species including fish and reptiles (69–71). It is expressed exclusively in the skeletal and cardiac muscles and is involved in myofibrillar assembly and force transmission in the heart (64), suggesting that the mechanism of Yakut cattle adaptation could be related to the function of the heart. In human and mice mutations in *NRAP* lead to dilated cardiomyopathy (73), a heart muscle genetic disorder, causing enlargement of ventricles and issues with pumping blood out of the heart. We hypothesise that the *NRAP* variant plays a crucial role in slowing the metabolism during the coldest periods in the Yakut cattle and other species, which would require enhanced heart ability to distribute blood at cold environmental temperatures. This is supported by strong enrichment for terms related to the cardiac contraction pathway in the Yakut cattle selected regions and the metabolic process and energy metabolism as well.

Scans for selection signatures pointed to multiple candidate genes and pathways for climate adaptations in Russian cattle breeds. A surprising result was that the Yakut cattle contains a large number of genes with signatures of negative selection and a very few with signatures of positive selection according to our dN/dS analysis. A possible explanation could be that the ancestral variants present in the Yakut cattle genome already make them a good fit into their environmental conditions, therefore a very few novel functional changes (like one in the *NRAP* gene) are required. Alternatively, it is possible that some additional functional changes could affect either regulatory gene regions or be represented by a different type of mutations not studied in this work, e.g. by indels. Genes with the highest dN/dS selection signals in the Yakut cattle included genes involved in the cytoskeleton stabilization which plays an important role in the hibernating mammals during return from torpor to euthermy (88), pointing to another possible common mechanism of reaction to extreme cold temperatures in the Yakut cattle and other animals. One of the top ranked regions under selection in the Yakut cattle contains the *PTN* a key gene in preserving insulin sensitivity, driving adipose tissue lipid turnover and the regulator of energy metabolism and thermogenesis (86). Additionally, we identified two candidate genes for cold/diet adaptation in native Siberian human populations (*PLA2G2A* and *ANGPTL8*) involved in lipid metabolism (87) in selected intervals of the Yakut cattle genome suggesting common mechanisms of adaptation to Siberian climates in mammals.

In conclusion, this work has revealed different evolutionary histories of two Russian northernmost cattle breeds, of which one is a typical taurine cattle and another contains multiple ancestral genetic variants, which allowed its adaptation to harsh conditions of living above the Polar Circle. Surprisingly, very few truly novel high-frequency genetic variants were required for this extreme adaptation, but the only detected missense change of this type represents a unique example of a young amino acid residue convergent change limited to a single cattle breed but shared with at least 16 species of hibernating/cold adapted mammals from nine distinct phylogenetic orders.

## Acknowledgements

This work was funded in part by the Russian Foundation for Basic Researches (RFBR) grant 18-016-00185 to DML. LB was funded by Marie Skłodowska-Curie grant agreement No. 703376. The identification and functional annotation of high-frequency breed-specific variants was supported by the Kurchatov Genomics Center of IC&G (075-15-2019-1662). The authors thank the participants of the 1000 Bull Genomes Project for providing sequence data, Michael Hiller and Emma Teeling for help with the 120-mammalian genome alignments.

