## Supplementary figures for "Demographic history, cold adaptation, and recent NRAP recurrent convergent evolution at amino acid residue 100 in the world northernmost cattle from Russia"

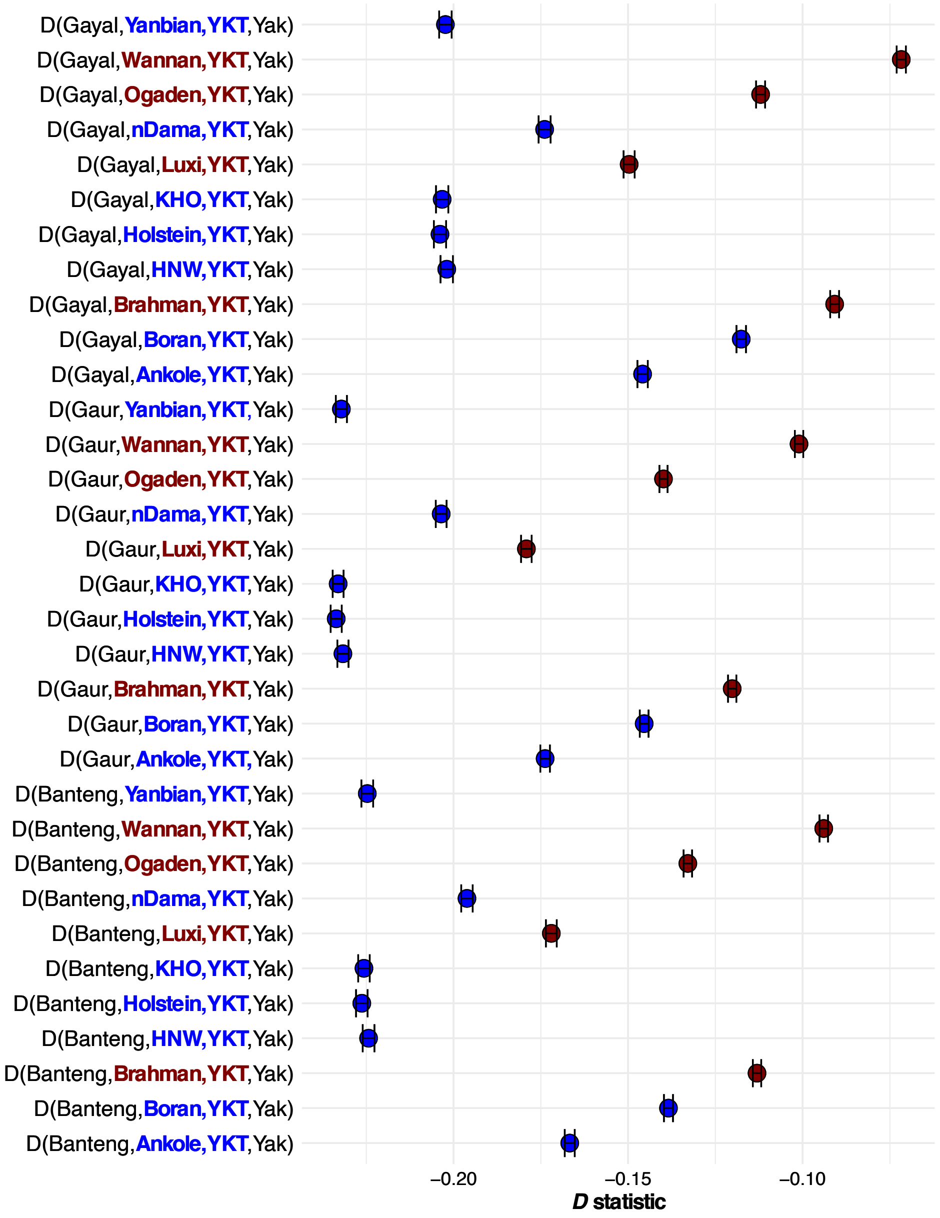


**Supplementary Figure 1.** D-statistics test, of the form D (A, B, X, Y), for admixture providing information about the direction of gene flow. Negative D-statistic indicates that gene flow occurred either between A and B or X and Y; cattle taurine breeds are in blue and cattle indicine breeds are in red. Abbreviations are as follow: YKT (Yakut), KHO (Kholmogory), HNW (Hanwoo).

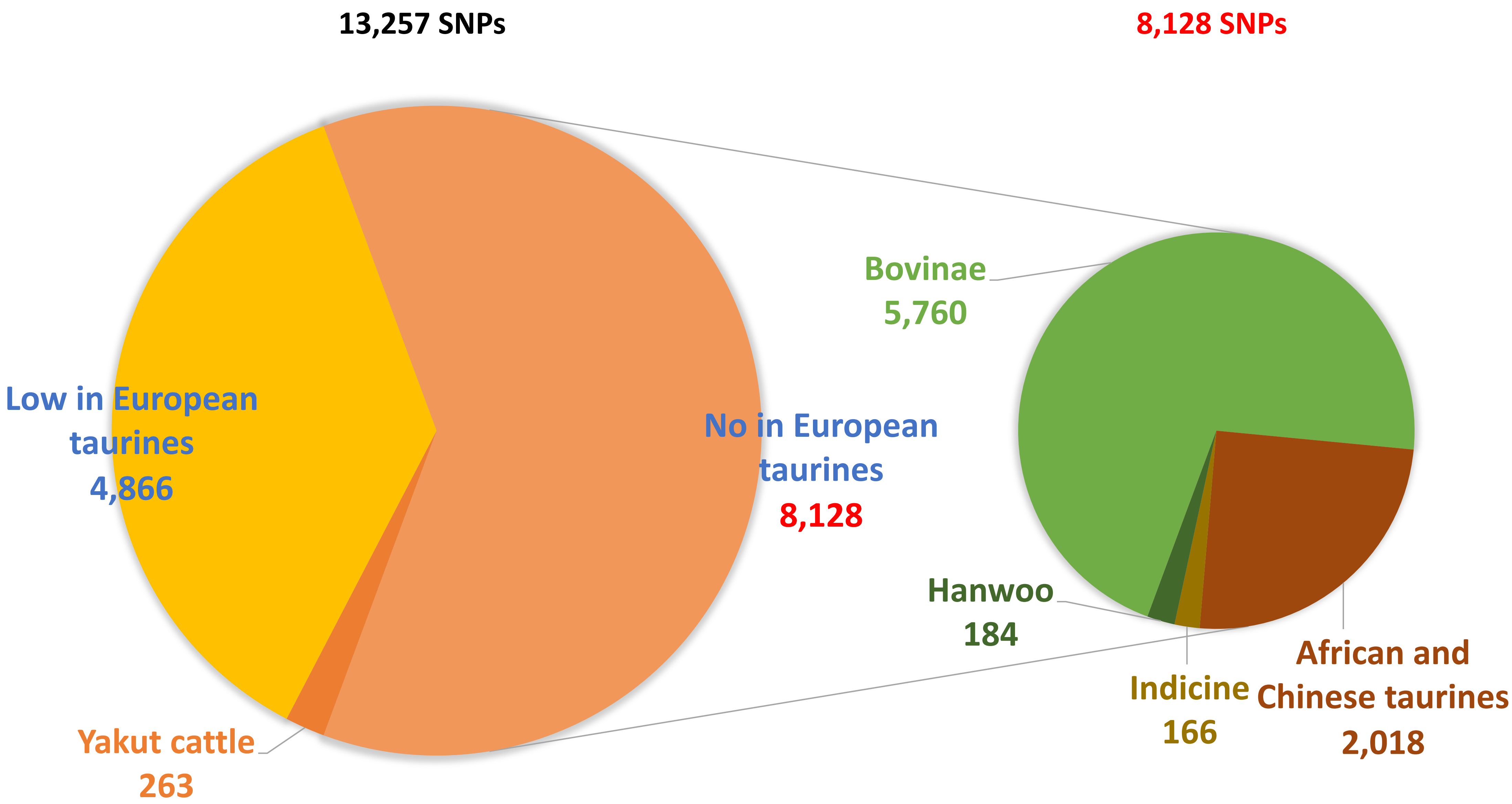


**Supplementary Figure 2.** Distribution of Yakut high frequency SNPs (total of 13,257 SNP) in European taurine, indicine cattle, Hanwoo, African, Indian, Chinese taurine, and bovinae species.

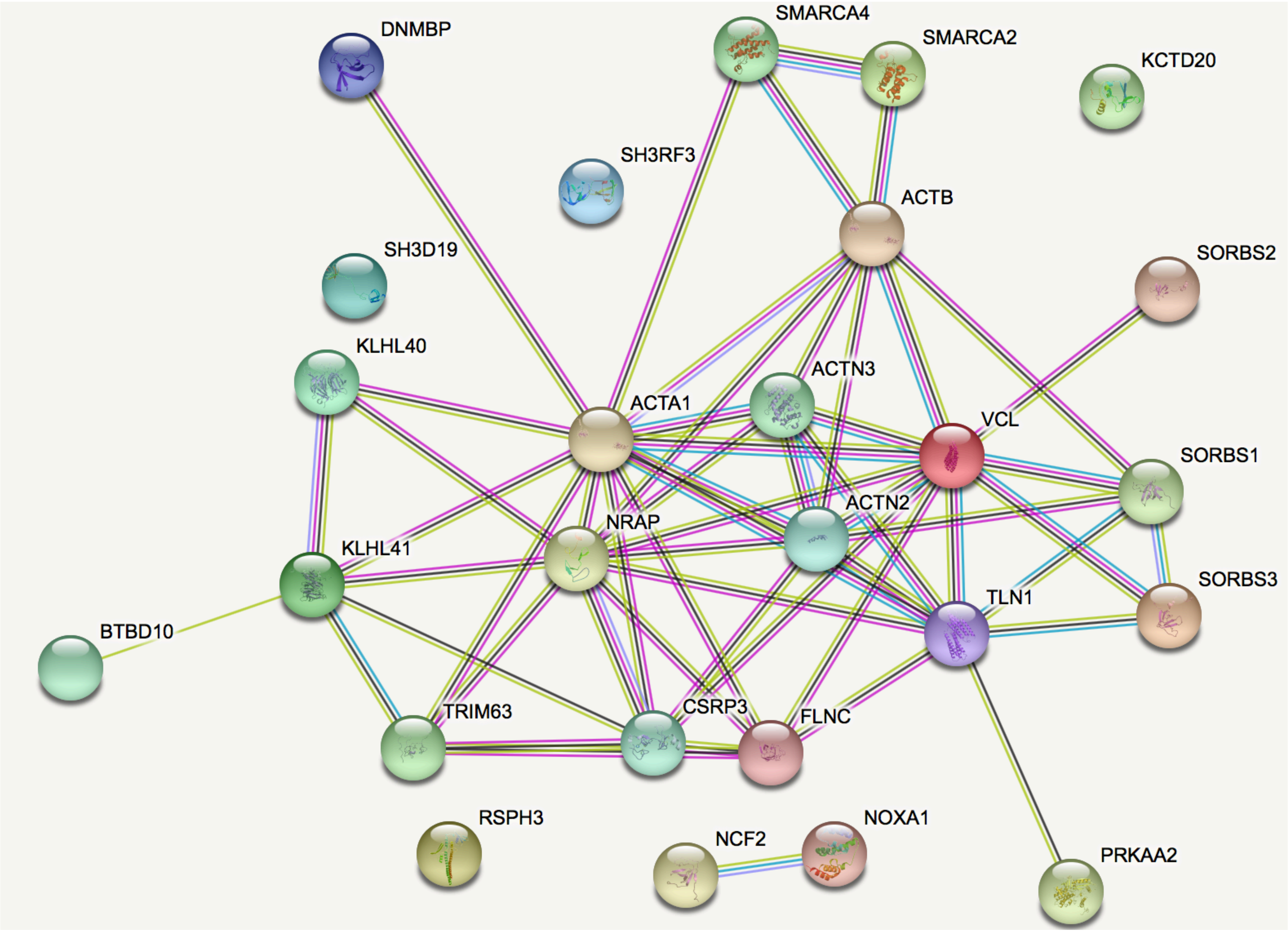


**Supplementary Figure 3.** ACTA1 gene network analysis.
